# Regulation of microtubule abundance and minus end dynamics by Katanin, CAMSAPs, WDR47 and kinesin-13

**DOI:** 10.64898/2026.03.26.714132

**Authors:** Dipti Rai, Elena Radul, Shasha Hua, Marèl F.M. Spoelstra, Eugene A. Katrukha, Kelly E. Stecker, Kai Jiang, Anna Akhmanova

## Abstract

Microtubule networks are major determinants of cell architecture and logistics. Microtubule organization and density are regulated by severing enzymes, which cut microtubule lattices or affect their growth and shortening. These activities can lead to microtubule amplification or disassembly, depending on the presence of microtubule stabilizers or destabilizers, but the interplay between these factors is poorly understood. Here, we reconstituted in vitro the activity of microtubule severase katanin together with microtubule minus-end stabilizers CAMSAPs, their binding partner WDR47 and microtubule depolymerase kinesin-13/MCAK. We confirmed that katanin can amplify or destroy microtubules in a concentration-dependent manner. CAMSAPs recruit katanin to microtubules and reduce katanin concentration needed for both amplification and destruction, whereas kinesin-13 completely abolishes microtubule amplification. WDR47 binds to microtubules decorated by CAMSAPs and suppresses katanin binding and severing. In addition, both katanin and WDR47 inhibit polymerization of CAMSAP-decorated microtubule minus ends. These data explain how these proteins act together to fine-tune microtubule minus-end stability without strongly increasing microtubule abundance.

Graphical abstract

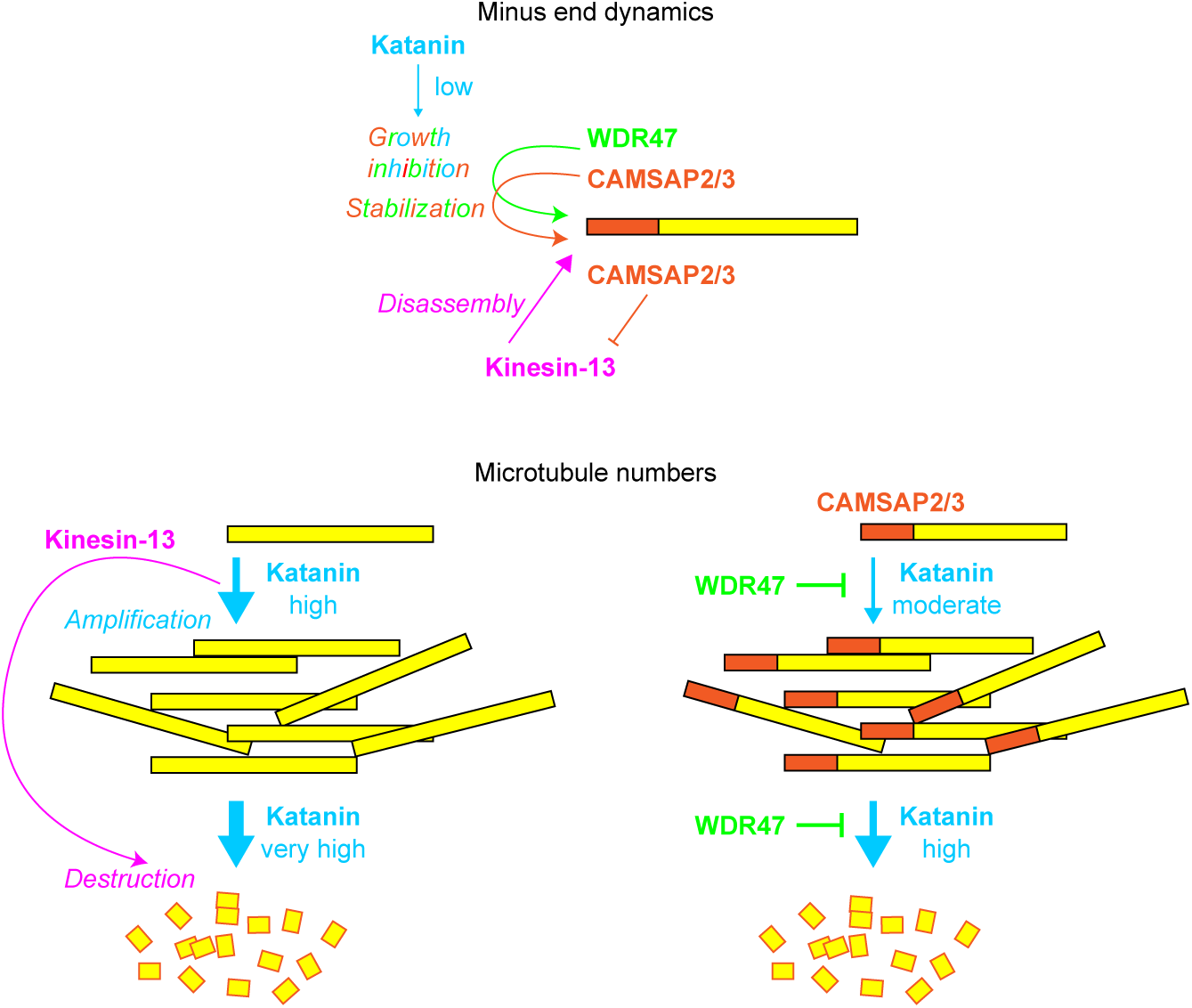

## Introduction

Microtubules are cytoskeletal filaments that serve as rails for intracellular transport, control organelle positioning, cell polarity, motility and chromosome separation during cell division. The density and geometry of microtubule arrays are critically important for cell architecture and function (Akhmanova and Kapitein, 2022; Burute and Kapitein, 2019; van Grinsven and Akhmanova, 2025). Microtubules are polarized polymers with two distinct ends – the slowly growing minus end and rapidly growing plus end (Akhmanova and Steinmetz, 2015; Desai and Mitchison, 1997). While polymerization of the plus ends generates most of the microtubule mass, the minus ends are often undynamic due to their anchoring at microtubule-organizing centers (MTOCs) (Sanchez and Feldman, 2017; Vineethakumari and Luders, 2022; Wu and Akhmanova, 2017). Microtubules are nucleated at MTOCs by the γ-tubulin ring complex (γ-TuRC), which can also cap the minus-ends (Liu et al., 2021b; Thawani and Petry, 2021; van Grinsven and Akhmanova, 2025). An alternative way of generating new microtubules is through cutting them into fragments by severing enzymes, such as katanin and spastin, and subsequent stabilization (Kuo and Howard, 2021; Vemu et al., 2018). Free, uncapped microtubule minus ends in animal cells can be decorated and stabilized by the members of CAMSAP/Patronin family (Hendershott and Vale, 2014; Jiang et al., 2014), reviewed in (Akhmanova and Steinmetz, 2019). Among the three mammalian CAMSAPs, CAMSAP1 tracks growing microtubule minus ends, whereas CAMSAP2 and CAMSAP3 stably decorate microtubule lattice grown from the minus end (Hendershott and Vale, 2014; Jiang et al., 2014). CAMSAP2 and 3 can also release microtubules from γ-TuRC (Rai et al., 2024). Microtubule stretches decorated by CAMSAP2/3 are very stable and can serve as “seeds” for microtubule re-growth (Jiang et al., 2014), providing a way to generate dense non-centrosomal microtubule arrays in large cells with extensive microtubule networks, such as epithelial cells and neurons (Dong et al., 2017; Khanal et al., 2016; King et al., 2014; Noordstra et al., 2016; Pongrakhananon et al., 2018; Toya et al., 2016; Yau et al., 2014; Zhou et al., 2020; Zhou et al., 2024).

CAMSAP-mediated microtubule minus-end protection is regulated in cells in two ways. First, microtubule depolymerases from the kinesin-13 family, KIF2A and KIF2C/MCAK, can disassemble microtubules and counteract CAMSAP-mediated microtubule stabilization (Atherton et al., 2017; Goodwin and Vale, 2010; Guan et al., 2023). Second, microtubule-severing ATPase katanin directly binds to CAMSAP2 and 3, the two stretch-forming CAMSAPs, and limits the length of microtubule segments decorated by these proteins (Jiang et al., 2018; Jiang et al., 2014). The biochemical interplay between CAMSAPs and katanin is further complicated by the demonstration that a WD40 repeat-containing protein WDR47 binds to CAMSAPs in the region close to the katanin-binding helix (**Fig. S1A-C**) (Buijs et al., 2021; Ren et al., 2022). WDR47 plays an essential role in controlling different aspects of brain and neuronal development (Buijs et al., 2021; Chen et al., 2020; Kannan et al., 2017), is recruited to cellular microtubules by CAMSAPs and promotes accumulation of CAMSAP2/3 on microtubules in neurons (Buijs et al., 2021; Chen et al., 2020). Experiments with overexpressed WDR47 and katanin in neurons led to the hypothesis that WDR47 may protect CAMSAP2/3-bound microtubule minus ends from katanin-mediated severing (Buijs et al., 2021). A related mechanism may operate during the biogenesis of motile cilia, where katanin, CAMSAPs and WDR47 cooperate in generating the central pair of axonemal microtubules (Liu et al., 2021a). The biochemical basis of the interplay between katanin, CAMSAPs, WDR47 and kinesin-13 is currently unclear. In vitro data demonstrate that severing enzymes tend to promote, rather than restrict the regrowth of microtubules they cut and thus increase microtubule numbers (Kuo et al., 2019; Vemu et al., 2018). If katanin amplifies microtubules, and CAMSAPs protect them, why does katanin then act as a negative regulator of CAMSAP-mediated microtubule-stabilization in cells? Is it due to the activity of kinesin-13 depolymerases, which use severase-generated damage as entry points for microtubule disassembly (Henkin et al., 2023)? Is WDR47 a competitive inhibitor CAMSAP-katanin interaction, or is another mechanism involved? How do the cellular concentrations of the studied factors compare to the concentrations needed to affect microtubule amplification or stability in vitro?

To address these questions, we reconstituted individual and collective activities of katanin, the three mammalian CAMSAPs, WDR47 and kinesin-13/KIF2C/MCAK in vitro on dynamic microtubules. We found that katanin could amplify or destroy dynamic microtubules in the presence of free tubulin, but only at high, likely non-physiological concentrations. CAMSAP2 and 3 recruited katanin to microtubules and lowered the concentration of katanin needed for both microtubule amplification and disassembly. The presence of kinesin-13 in combination with katanin dramatically shifted the balance towards microtubule depolymerization, explaining why in many cellular settings, katanin is a negative and not a positive regulator of microtubule density, as one could assume from the in vitro work. WDR47 protected CAMSAP2/3 decorated microtubules by directly inhibiting katanin binding, explaining the activity of this protein in generation of stable microtubule arrays in neurons and cilia. Finally, all studied proteins limited the ability of microtubule minus ends to grow, explaining why minus-end polymerization, which is readily detectable with purified tubulin, is hard to detect even in cells with many uncapped minus ends.

## Results

### Katanin amplifies or destroys microtubules in a concentration-dependent manner

To study the activities of different microtubule regulators, we purified them from transiently transfected HEK293 cells using Twin-Strep-tag as described previously (Jiang et al., 2017; Rai et al., 2024). We performed in vitro reconstitution assays with microtubules grown from GMPCPP-stabilized seeds and observed microtubule growth, amplification or disassembly by Total Internal Reflection Fluorescence (TIRF) microscopy (Bieling et al., 2007; Jiang et al., 2017). Since katanin can rapidly alter microtubule numbers, we first allowed microtubules to grow from immobilized seeds and then flowed in different concentrations of katanin **(Fig. 1A)** and quantified microtubule abundance using Fiji plugin FeatureJ: Hessian (see Methods for details). Our previous work showed that the complex of fluorescently tagged katanin p60 subunit and the C-terminal part of the p80 subunit lacking the WD40 domain (p60/p80C) **(Fig. S1A, D)** can cut dynamic microtubules in vitro in the presence of free tubulin, but severing required concentrations in the several-hundred-nanomolar range (Jiang et al., 2017). We confirmed these observations: at low katanin concentrations (e.g. 40 nM, **Fig. 1B**), p60/p80C did not bind or cut dynamic microtubules, likely because free tubulin inhibits katanin activity (Bailey et al., 2015). At 100 nM, a concentration close to the physiological katanin concentration, which was estimated to be 60-100 nM in HeLa and HEK293 cells (Cho et al., 2022; Nagaraj et al., 2011), occasional microtubule cutting was observed **(Fig. 1B, C** and **Video S1)**. Abundant severing and strong microtubule amplification were observed with 200 nM p60/p80C, whereas at 400 nM, all microtubules were destroyed **(Fig. 1D-F** and **Video S1)**. For comparison, we prepared a fluorescently tagged full-length katanin containing not only the whole p60 subunit but also the complete p80 subunit, including the WD40 domain (p60/p80) **(Fig. S1A, D)**. Analysis of this complex by mass spectrometry revealed no significant contamination with any microtubule-binding proteins **(Fig. S1E** and **Table S1)**. Full-length p60/p80 was less active than p60/p80C and showed very little severing of dynamic microtubules even at 250 nM **(Fig. S1F)**, the highest concentration we could test with this purified protein due to its tendency to aggregate.

**Figure 1.**
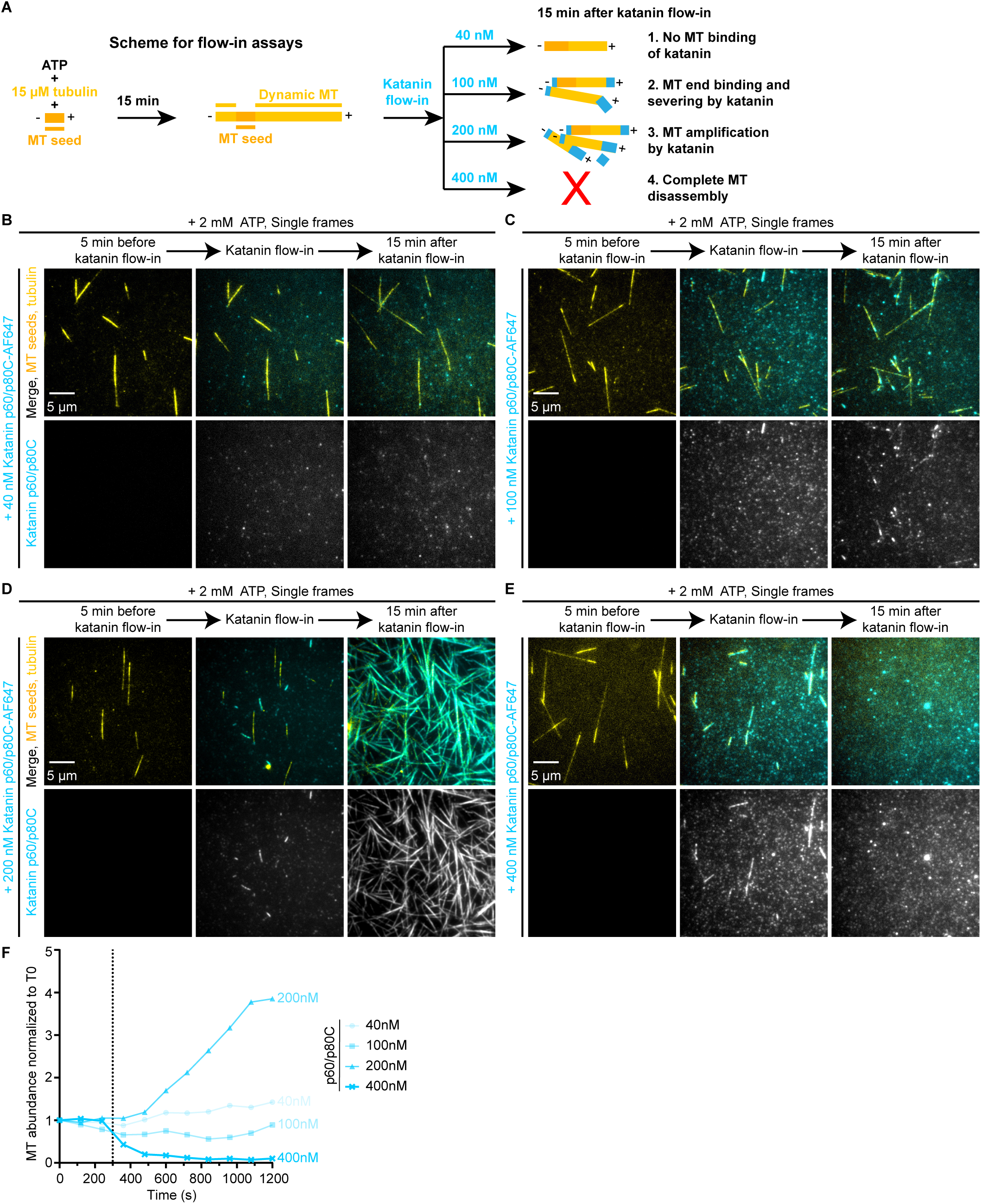
Katanin p60/p80C amplifies or destroys microtubules in a concentration-dependent manner. **(A)** A scheme for in vitro reconstitution of microtubule severing by katanin. Microtubules were polymerized for 15 min before the substitution of microtubule polymerization mix with severing mix (same as the polymerization mix but supplemented with katanin) during flow-in. Katanin activity was monitored for 15 min after flow-in. **(B-E)** Single frames (at indicated steps) from 20 min time-lapse movies showing katanin (cyan) mediated regulation of microtubule (yellow) amplification and destruction in the presence of different concentrations of katanin p60/p80C-SNAP-AF647, ranging from low to very high: 40 nM **(B)**, 100 nM **(C)**, 200 nM **(D),** and 400 nM **(E)**, under indicated assay conditions, also depicted in panel **(A)**. **(F)** Quantification of microtubule abundance over time normalized to microtubule abundance at t=0 min (T0 being 5 min before katanin flow-in) in the presence of different concentrations of katanin p60/p80C-SNAP-AF647 alone (as indicated on plot) under the assay conditions represented in panels **(B-E)**. n=3, N=3 for each condition, where n is the total number of single fields of view analyzed from N number of independent assays. Each individual curve represents the mean value of the three fields of view analyzed per condition.

### Katanin inhibits growth, severs and amplifies CAMSAP2/3 decorated microtubules

We next tested whether CAMSAPs potentiate katanin-mediated microtubule severing. We purified three mammalian CAMSAP isoforms **(Fig. S2A)**, as described previously (Jiang et al., 2014; Rai et al., 2024). Microtubules were grown in the presence of one of the CAMSAPs for 15 min, followed by flow-in of a mix of the same CAMSAP and different concentrations of p60/p80C **(Fig. 2A)**. In the presence of 40 nM CAMSAP3 (a concentration similar to the physiological CAMSAP3 concentration measured in HEK293 cells (Cho et al., 2022)), p60/p80C was clearly detectable at microtubule minus ends already at 5 nM **(Fig. 2B, C** and **Video S2)**. Binding of p60/p80C inhibited microtubule minus-end polymerization **(Fig. 2D)**, similar to the previously observed effect of the katanin-ASPM combination (Jiang et al., 2017). When the concentration of p60/p80C was elevated to 40 nM, it triggered rapid and specific fragmentation of CAMSAP3-decorated microtubule stretches **(Fig. 2E** and **Video S2)**. The resulting fragments persisted and initiated growth of new microtubules, leading to the formation of bundled brush-like structures and a strong overall increase of microtubule numbers **(Fig. 2E, F, H)**. A similar effect was observed when 40 nM p60/p80C was added to microtubules decorated by 40 nM CAMSAP2, although the cutting efficiency and the ensuing microtubule amplification were somewhat lower than with CAMSAP3 **(Fig. 2F** and **Fig. S2B, C)**. In contrast, CAMSAP1, which does not interact with katanin (Jiang et al., 2014), did not recruit p60/p80C to microtubules and did not potentiate their severing or amplification **(Fig. 2F** and **Fig. S2C, D)**.

**Figure 2.**
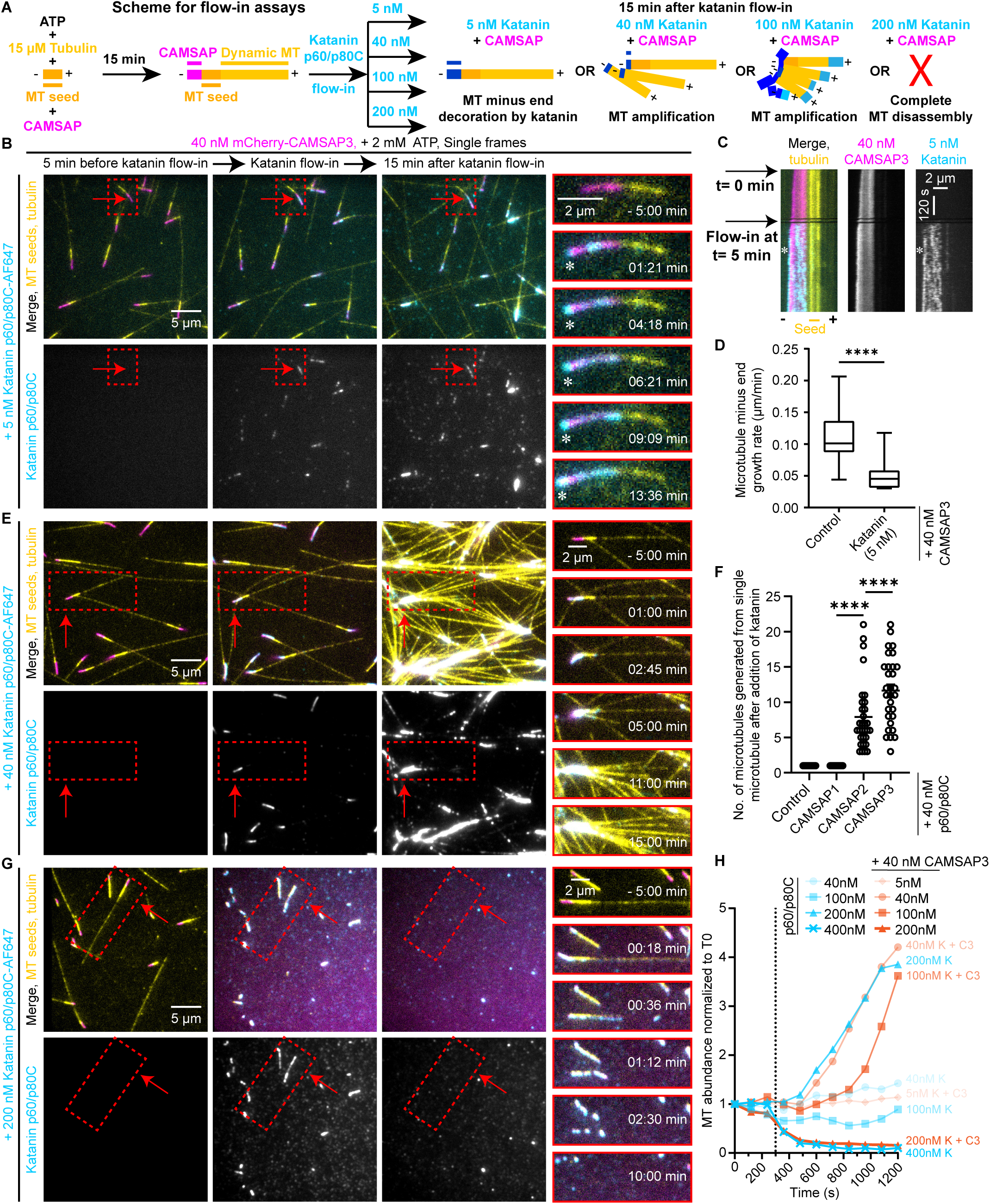
Katanin p60/p80C inhibits minus end growth of CAMSAP-decorated microtubules and regulates microtubule abundance by severing them. **(A)** A scheme for in vitro reconstitution of CAMSAP-decorated microtubule minus end regulation by katanin. **(B, E, G)** Single frames (at indicated steps) from 20 min time-lapse movies showing regulation of minus end dynamics **(B)**, severing mediated amplification **(E)** and complete disassembly **(G)** of mCherry-CAMSAP3-decorated (magenta) microtubules (yellow) by 5 nM **(B)**, 40 nM **(E)**, and 200 nM **(G)** katanin p60/p80C-SNAP-AF647 (cyan), respectively, under indicated assay conditions, also depicted in panel **(A)**. The magnified still frames (with -05:00 min being 5 min before katanin flow-in) on right show katanin binding and its subsequent severing activity on individual CAMSAP-decorated microtubules, highlighted by red boxes and arrows. White asterisks in the magnified views show katanin binding to the outmost minus end of the microtubule. **(C)** Representative kymographs from the in vitro reconstitution experiments shown in panel **B**. **(D)** The box and whisker plot showing average growth rate of microtubule minus ends in the presence of 40 nM mCherry-CAMSAP3 with (n=35, m=35, N=4) or without (n=41, m=41, N=4) 5 nM katanin p60/p80C, quantified from the assays represented in panel **(B, C)**. n is the number of growth events; m is the number of microtubules analyzed from N number of independent experiments for each condition. The box extends from the 25th to 75th percentiles, while the whiskers extend to the smallest and largest values. The line in the middle of the box is the median. Two-tailed unpaired t-test was used to test for significance. ****p< 0.0001. **(F)** Quantification of the number of microtubules generated from a single CAMSAP-decorated microtubule as a result of katanin-mediated severing under the experimental conditions shown in panel **(E)**, Fig. 1B and **Fig. S2B, D**. n=47, N=3 for katanin p60/p80C-SNAP-AF647 alone; n=49, N=3 for CAMSAP1; n=31, N=3 for CAMSAP2 and n=30, N=3 for CAMSAP3, where n is the number of single microtubules analyzed and N is the number of independent assays. The plot presents mean ± s.e.m., and individual data points represent single microtubules analyzed. One-way ANOVA with Tukey’s multiple comparisons test was performed to compare the means with each other (****p< 0.0001). **(H)** Quantification of microtubule abundance over time normalized to microtubule abundance at t=0 min (T0 being 5 min before katanin flow-in) in the presence of different concentrations of katanin p60/p80C-SNAP-AF647 (as indicated on plot) either alone or together with 40 nM mCherry-CAMSAP3 under the assay conditions represented in panels **(B, E, G)**, Fig. 1B**-E** and **Fig. S2E**. n=3, N=3 for each condition, where n is the total number of single fields of view analyzed from N number of independent assays. Each individual curve represents the mean value of the three fields of view analyzed per condition. Data for different concentrations of katanin p60/p80C alone are from Fig. 1F, replotted here for comparison.

When we increased the concentration of p60/p80C to 100 nM, fragmentation of CAMSAP3-decorated microtubule stretches became even stronger, while microtubule outgrowth from these stretches was suppressed, and at 200 nM, p60/p80C completely disassembled all microtubules decorated by 40 nM CAMSAP3 **(Fig. 2G, H, Fig. S2E** and **Video S2)**.

We also performed similar experiments with full-length p60/p80 **(Fig. 3A)**. It was less active than the p60/p80C version, because at 40 nM, it did not sever microtubules decorated with 40 nM CAMSAP3, although it did sever CAMSAP3-bound microtubules and amplified microtubule numbers when added at a 250 nM concentration **(Fig. 3B-E** and **Video S3)**. It should be noted that in these conditions, the actual concentration of katanin in solution could be lower, because we observed significant absorption of katanin on the coverslip surface **(Fig. 3D)**. We also combined 250 nM full-length katanin with microtubules decorated with CAMSAP1 and CAMSAP2 but observed little microtubule binding or severing in these conditions **(Fig. S3A, B)**. This is in line with the observation that CAMSAP2 potentiated p60/p80C-mediated microtubule severing less efficiently than CAMSAP3, and CAMSAP1 did not promote microtubule severing at all **(Fig. 2F** and **Fig. S2C)**. The inability of full-length katanin to efficiently sever CAMSAP2-bound microtubules could be due to a lower affinity of katanin for CAMSAP2, or a weaker microtubule decoration by CAMSAP2 compared to CAMSAP3 as observed previously (Jiang et al., 2014). We conclude that CAMSAP2-, and particularly CAMSAP3-decorated microtubules can serve as a specific substrate for katanin-mediated microtubule severing, reducing the concentration of katanin needed both for microtubule amplification and destruction.

**Figure 3.**
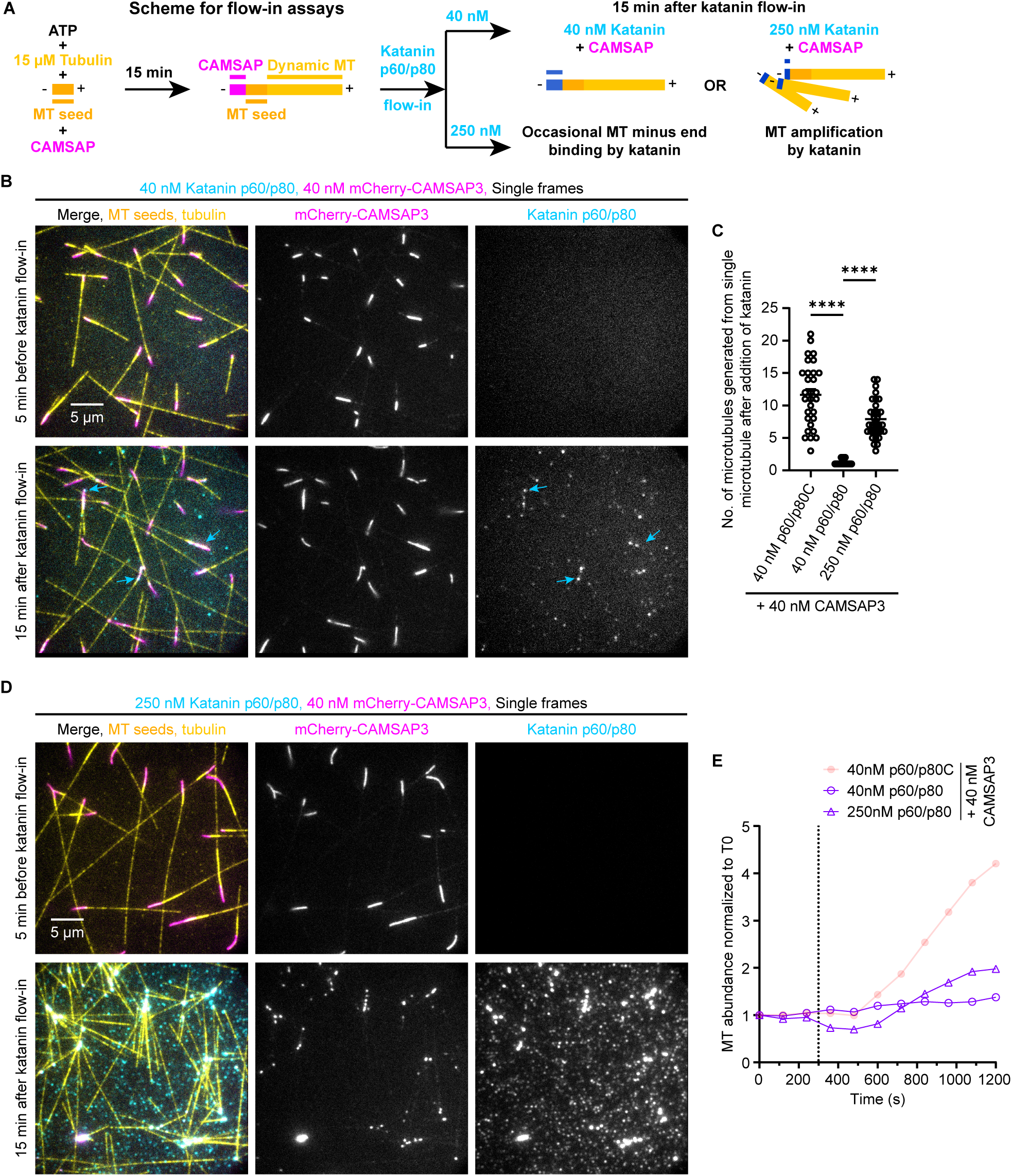
Full-length katanin p60/p80 severs and amplifies CAMSAP3-decorated microtubules. **(A)** A scheme for in vitro reconstitution of microtubule severing by full-length katanin. **(B, D)** Single frames (at indicated steps) from 20 min time-lapse movies showing katanin (cyan) binding and its subsequent severing activity on mCherry-CAMSAP3-decorated (magenta) microtubules (yellow) at 40 nM **(B)** or 250 nM **(D)** concentrations of full-length katanin p60/p80-SNAP-AF647, under indicated assay conditions, also depicted in panel **(A)**. Cyan arrows denote the occasional binding of full-length katanin to CAMSAP3-decorated minus ends at 40 nM katanin concentration. **(C)** Quantification of the number of microtubules generated from a single CAMSAP3-decorated microtubule as a result of katanin-mediated severing under the assay conditions represented in panels **(B, D)** and Fig. 2E. n=30, N=3 for 40 nM p60/p80C; n=58, N=3 for 40 nM p60/p80 and n=31, N=3 for 250 nM p60/p80, where n is the number of single microtubules analyzed from N number of independent assays. The plot presents mean ± s.e.m. and individual data points represent single microtubules analyzed. One-way ANOVA with Šídák’s multiple comparisons test was performed for the pairwise comparison of means. ****p< 0.0001. Data for 40 nM p60/p80C with CAMSAP3 are from Fig. 2F, replotted here for direct comparison. **(E)** Quantification of microtubule abundance over time normalized to microtubule abundance at t=0 min (T0 being 5 min before katanin flow-in) in the presence of 40 nM or 250 nM full-length katanin p60/p80-SNAP-AF647 (as indicated on plot) together with 40 nM mCherry-CAMSAP3 under the assay conditions represented in panels **(B, D)** and Fig. 2E. n=3, N=3 for each condition, where n is the total number of single fields of view analyzed from N number of independent assays. Each individual curve represents the mean value of the three fields of view analyzed per condition. Data for 40 nM p60/p80C with CAMSAP3 are from Fig. 2F, replotted here for direct comparison.

### Kinesin-13 potently promotes depolymerization of microtubules severed by katanin

Cells express two major microtubule depolymerases from kinesin-13 family, which can disassemble microtubule plus- and minus ends (Walczak et al., 2013). Here, we used a preparation of kinesin-13 KIF2C/MCAK **(Fig. S4A)**. Mass spectrometry analysis showed mild (a few percent) contamination with the binding partners of MCAK, CEP170 and EB1 (Lee et al., 2008; Welburn and Cheeseman, 2012), which could potentially enhance its localization to microtubule ends **(Fig. S4B** and **Table S2)**. At 5 nM, mCherry-MCAK did not block microtubule growth but promoted catastrophes, as expected (Montenegro Gouveia et al., 2010) **(Fig. 4A-C** and **Video S4)**. However, when combined with 100 nM p60/p80C, a katanin concentration sufficient to induce only occasional cutting, 5 nM MCAK promptly triggered complete microtubule disassembly **(Fig. 4D, E** and **Video S4)**. This is probably due to the fact that not only the microtubules that were severed but also those that were partially damaged by katanin and could be subsequently repaired (Vemu et al., 2018), were now depolymerized. Cellular MCAK concentrations have been estimated in the range of 10 to 250 nM in different cell types (Cho et al., 2022; Nagaraj et al., 2011), higher than the one used in our assays. This indicates that kinesin-13 can powerfully suppress katanin-mediated microtubule amplification. To confirm this idea, we combined 5 nM mCherry-MCAK with 40 nM GFP-CAMSAP3 and then flowed in 100 nM katanin in the presence of the same proteins **(Fig. 4F)**. CAMSAP3-decorated microtubule stretches were protected from MCAK-induced disassembly **(Fig. 4G, H** and **Video S4)**. However, in contrast to the conditions without MCAK **(Fig. S2E** with mCherry-CAMSAP3 and **Fig. S4C** with GFP-CAMSAP3), where we observed massive microtubule amplification, only a limited increase in microtubule abundance was observed **(Fig. 4I, J** and **Video S4)**. Remarkably, microtubule numbers were almost maintained in a steady state when MCAK was combined with 40 nM katanin **(Fig. S4D)**. These data help to explain why in cells, katanin antagonizes formation of long CAMSAP-decorated stretches without causing a strong increase in microtubule density and acts as a negative regulator of microtubule stability (Ahmad et al., 1999; Buijs et al., 2021; Jiang et al., 2018; Jiang et al., 2014; Nieuwenhuis et al., 2017).

**Figure 4.**
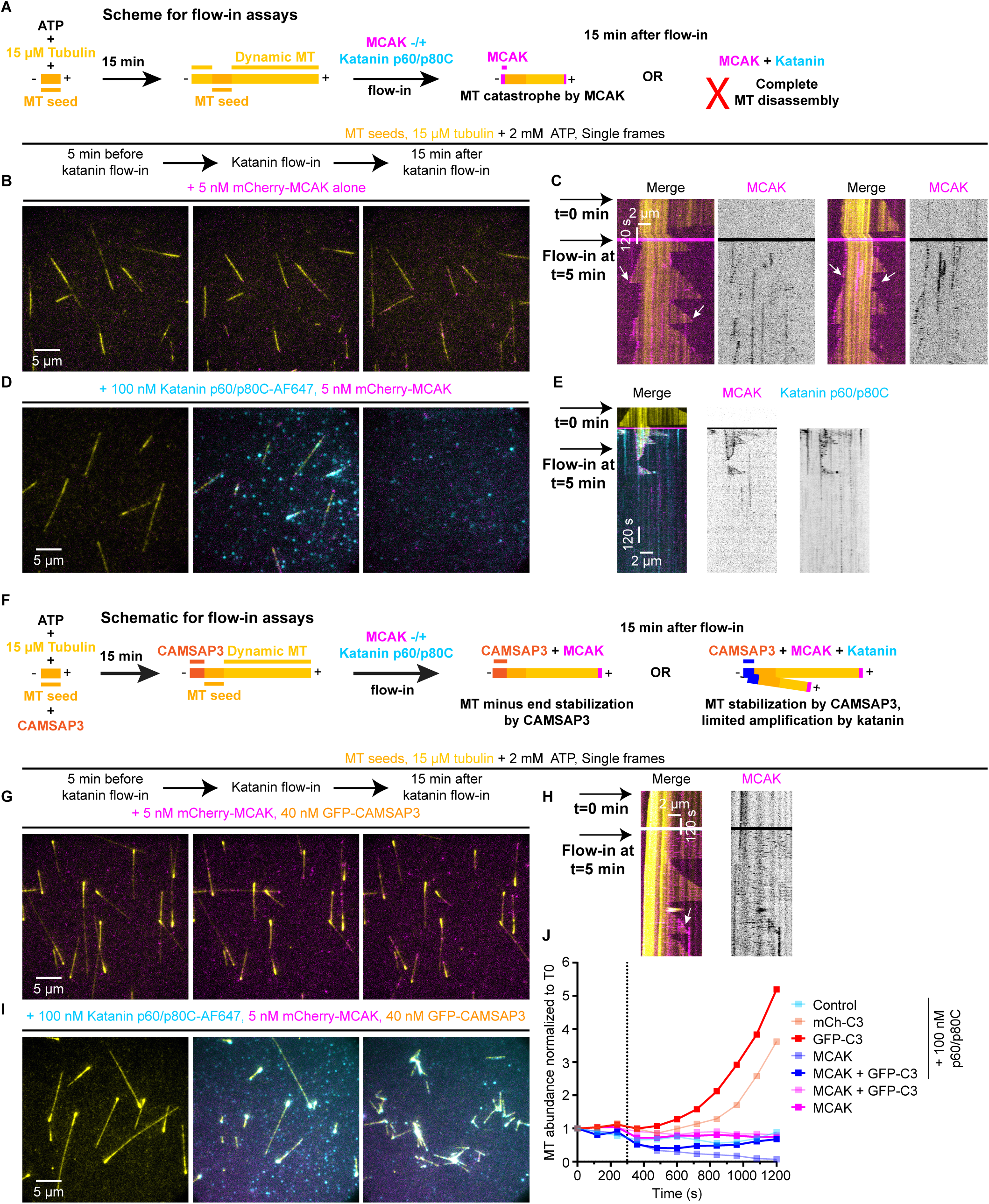
Kinesin-13 potently promotes depolymerization of microtubules severed by katanin, while CAMSAP3 helps to maintain a steady state of microtubule numbers. **(A)** A scheme for in vitro reconstitution of microtubule severing by katanin and its regulation by MCAK/kinesin-13. **(B, D)** Single frames (at indicated steps) from 20 min time-lapse movies showing depolymerization of microtubule (yellow) ends by 5 nM mCherry-MCAK alone (magenta) **(B)** and microtubule destruction by MCAK-mediated depolymerization and katanin-mediated severing by 5 nM mCherry-MCAK together with 100 nM katanin p60/p80C-SNAP-AF647 (cyan) **(D)** under indicated assay conditions, also depicted in panel **(A)**. **(C, E)** Representative kymographs from the in vitro reconstitution experiments represented in panel **B** for MCAK alone **(C)** and panel **D** for MCAK together with katanin **(E)**. White arrows show microtubule catastrophes. **(F)** A scheme for in vitro reconstitution of microtubule severing by katanin and its regulation by MCAK and CAMSAP3. **(G, I)** Single frames (at indicated steps) from 20 min time-lapse movies showing microtubule (yellow) minus-end stabilization by 40 nM GFP-CAMSAP3 (yellow) against depolymerization by 5 nM mCherry-MCAK (magenta) **(G)** and antagonizing activity of 40 nM GFP-CAMSAP3 against microtubule destruction by 5 nM mCherry-MCAK together with 100 nM katanin p60/p80C-SNAP-AF647 (cyan) **(I)** under indicated assay conditions, also depicted in panel **F**. **(H)** Representative kymographs from the in vitro reconstitution experiments represented in panel **G**. White arrow shows microtubule catastrophe at the plus end. **(J)** Quantification of microtubule abundance over time normalized to microtubule abundance at t=0 min (T0 being 5 min before flow-in) under the assay conditions indicated on plot, and represented in panels **(B, D, G, I)**, Fig. 1C**, Fig. S2E** and **Fig. S4C**. n=3, N=3 for each condition, where n is the total number of single fields of view analyzed from N number of independent assays. Each individual curve represents the mean value of the three fields of view analyzed per condition. Data for these assay conditions are reused from the following figure panels and replotted here for a direct comparison: 100 nM katanin p60/p80C alone from Fig. 1F; 100 nM katanin p60/p80C together with 40 nM mCherry-CAMSAP3 from Fig. 2H, respectively.

### WDR47 binds to CAMSAP-decorated minus ends and affects their dynamics

CAMSAP2 and 3 strongly recruit katanin to the microtubules they decorate, resulting in their fragmentation, but in cells, katanin also acts at other microtubule sites (Liu et al., 2021a; Yu et al., 2008), whereas CAMSAP-decorated microtubule ends remain stable. A potential factor that can limit katanin-mediated destruction of CAMSAP-decorated microtubules is WDR47 (Buijs et al., 2021; Chen et al., 2020). We used HEK293T cells to purify Twin-Strep- and GFP-tagged WDR47 **(Fig. S5A)**. Mass spectrometry-based analysis showed that this WDR47 preparation contained low levels of all three mammalian CAMSAPs, with CAMSAP1 being the most abundant **(Fig. S5B** and **Table S3)**. When included in the assay with dynamic microtubules at 15 nM concentration, WDR47 very occasionally bound to microtubule minus-ends **(Fig. 5A, B)**. However, minus end binding could be observed much more frequently in the presence of 150 nM WDR47 **(Fig. 5C)**. In such cases, WDR47 could block minus end growth (for a short time or for the whole duration of a 10-minute movie) or track growing minus ends without having a major effect on their growth rate **(Fig. 5C-E)**.

**Figure 5.**
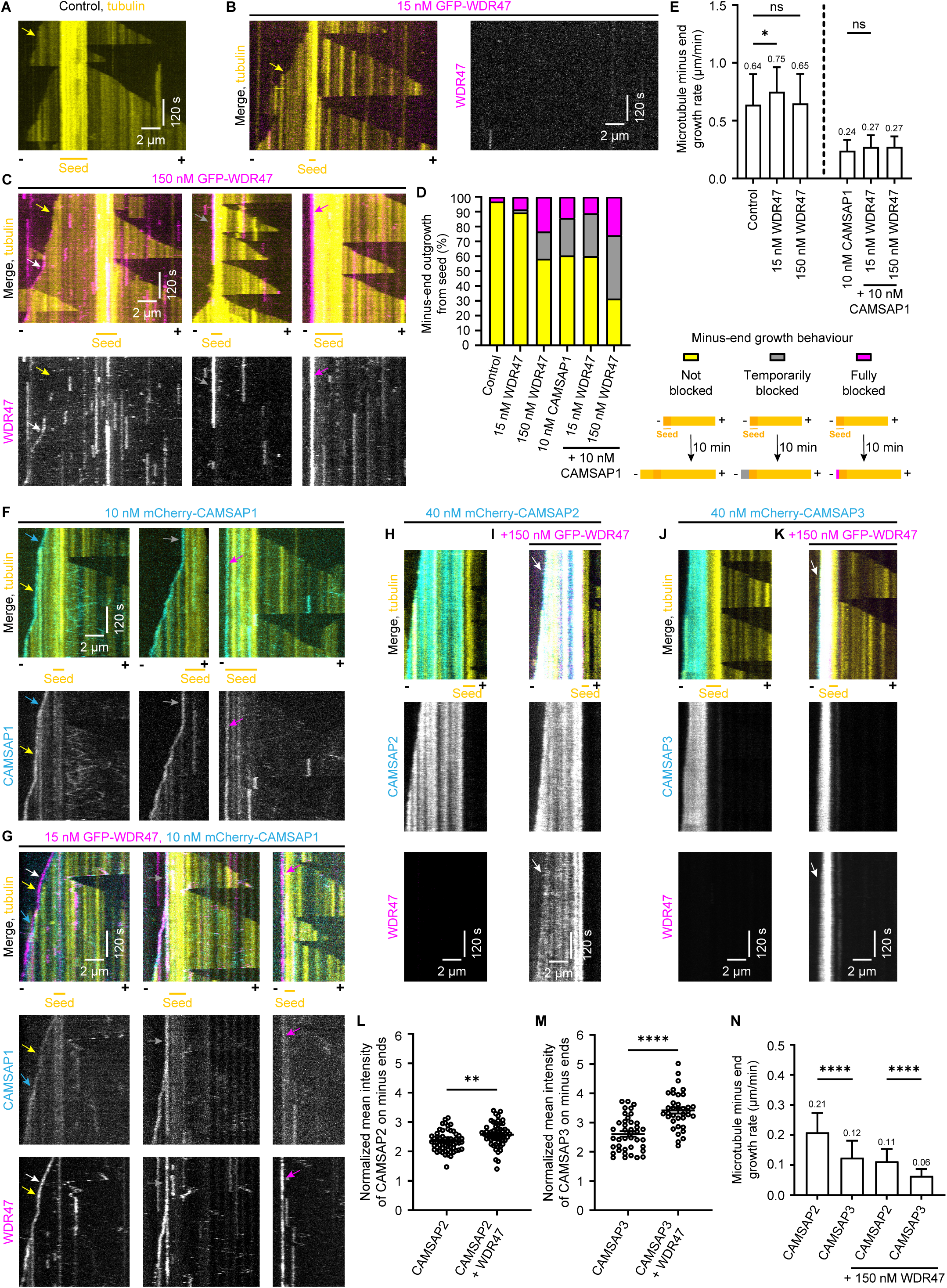
WDR47 binds to CAMSAP-decorated microtubule minus ends and affects their dynamics. **(A-C, F, G)** Representative kymographs from in vitro reconstitutions of microtubule (yellow) dynamics in the presence of either 15 μM tubulin (yellow) alone (**A**) or together with either 15 nM GFP-WDR47 (magenta) (**B**), or 150 nM GFP-WDR47 (**C**), or 10 nM mCherry-CAMSAP1 (cyan) **(F)**, or 10 nM mCherry-CAMSAP1 and 15 nM GFP-WDR47 **(G)**. In panels **(C, F, G)**, three different types of minus-end growth behaviors are illustrated (also depicted in the key for panel **D**): minus-end growth (left, yellow arrows); transition from temporarily blocked minus-end outgrowth to growing minus-end (middle and gray arrows) and completely blocked minus-end of a GMPCPP-stabilized microtubule seed (right and magenta arrows). White and cyan arrows denote tracking of minus ends by WDR47 and CAMSAP1, respectively. See also **Fig. S5C**. **(D)** Quantification of different types of minus-end growth behavior as described for panels **(A-C, F, G)** in the presence of 15 μM tubulin alone (n=94, N=4) or together with either 15 nM GFP-WDR47 (n=47, N=3), or 150 nM GFP-WDR47 (n=60, N=3), or 10 nM CAMSAP1 (n=91, N=5), or 10 nM CAMSAP1 and 15 nM GFP-WDR47 (n=45, N=3), or 10 nM CAMSAP1 and 150 nM GFP-WDR47 and (n=89, N=5); where n is the number of minus ends analyzed and N is the number of independent experiments analyzed for each condition. Representative kymographs are shown in panels **A-C, F, G,** and **Fig. S5C**. **(E)** Quantification of average growth rate of microtubule minus ends in the presence of 15 μM tubulin alone (n=111, N=3) or together with either 15 nM GFP-WDR47 (n=43, N=3), or 150 nM GFP-WDR47 (n=46, N=3), or 10 nM CAMSAP1 (n=48, N=3), or 10 nM CAMSAP1 and 15 nM GFP-WDR47 (n=34, N=3), or 10 nM CAMSAP1 and 150 nM GFP-WDR47 (n=41, N=4), where n is the number of growth events analyzed and N is the number of independent experiments analyzed for each condition. The plot presents mean ± s.d. and individual data points represent growth events analyzed. One-way ANOVA with Tukey’s multiple comparisons corrected for multiple testing was performed to compare the means with each other. not significant (ns), p=0.9997 between control and 150 nM WDR47; ns, p=0.9815 between 10 nM CAMSAP1 with or without 150 nM WDR47; *p=0.0305 between control and 15 nM WDR47. Representative kymographs are shown in panels **A-C, F-G,** and **Fig. S5C**. **(H-K)** Representative kymographs illustrating WDR47 (magenta) decoration of microtubule (yellow) minus ends stabilized by CAMSAPs (cyan) in the presence of 15 μM tubulin together with either 40 nM mCherry-CAMSAP2 **(H)**, or 40 nM mCherry-CAMSAP2 and 150 nM GFP-WDR47 **(I)**, or 40 nM mCherry-CAMSAP3 **(J)**, or 40 nM mCherry-CAMSAP3 and 150 nM GFP-WDR47 **(K)**. White arrows denote WDR47 decoration on CAMSAP2/3 stretches. **(L, M)** Plots showing normalized mean intensity of CAMSAP2 **(L)** and CAMSAP3 **(M)** stretches quantified (see method section for details) from the assay conditions indicated on the plots and represented in panels **(H-K)**. In panel **L**:15 μM tubulin together with either 40 nM CAMSAP2 (n=60, N=3) or 40 nM CAMSAP2 and 150 nM GFP-WDR47 (n=57, N=3). In panel **M**:15 μM tubulin together with either 40 nM CAMSAP3 (n=40, N=3) or 40 nM CAMSAP3 and 150 nM GFP-WDR47 (n=36, N=3). n is the number of minus ends analyzed, and N is the number of independent experiments for each condition. The plot presents mean ± s.e.m. and individual data points represent each minus end analyzed. Two-tailed unpaired t-test was used for significance test. **p= 0.0012; ****p< 0.0001. See also **Fig. S5D, E**. **(N)** Quantification of average growth rate of microtubule minus ends in the presence of 15 μM tubulin together with either 40 nM CAMSAP2 (n=45, N=3), or 40 nM CAMSAP3 (n=41, N=4), or 40 nM CAMSAP2 and 150 nM GFP-WDR47 (n=35, N=3), or 40 nM CAMSAP3 and 150 nM GFP-WDR47 (n=61, N=4); where n is the number of growth events analyzed and N is the number of independent experiments analyzed for each condition. The plot presents mean ± s.d. and individual data points represent growth events analyzed. One-way ANOVA with Tukey’s multiple comparisons corrected for multiple testing was performed to compare means with each other. ****p< 0.0001. Representative kymographs are shown in panels (**H-K)**.

Since minus end binding and tracking are behaviors typical for CAMSAP1, we hypothesized that they were caused by CAMSAP1 contamination in our WDR47 preparations. In agreement with this idea, assays with 10 nM CAMSAP1 alone showed that it was present at the minus ends of most microtubules, could block minus-end outgrowth from the seeds or track growing microtubule minus ends; transitions from blocked to growing minus ends were also observed **(Fig. 5D, F)**. To get further support for the idea that WDR47 was targeted to the minus ends by CAMSAP1 present in the preparation, we combined 15 nM WDR47 (a concentration at which it did not show any significant minus end binding) with 10 nM CAMSAP1 and observed that the two proteins colocalized on either growing or blocked microtubule minus ends **(Fig. 5G)**. Similar minus end binding and blocking were observed when 150 nM WDR47 was combined with 10 nM CAMSAP1, but increasing WDR47 concentration in the presence of CAMSAP1 made minus-end blocking more frequent **(Fig. 5D** and **Fig. S5C)**. CAMSAP1 by itself slowed down the growth rate of microtubule minus ends compared to control, and this effect remained unaltered by the addition of WDR47 **(Fig. 5E)**.

We also combined WDR47 with CAMSAP2 or CAMSAP3, which decorate microtubule lattice grown from the minus end, slow down microtubule minus-end polymerization and prevent catastrophes at the minus end (Jiang et al., 2014). When 150 nM WDR47 was added to microtubules in the presence of 40 nM CAMSAP2 or CAMSAP3, it strongly decorated the whole microtubule minus-end-grown part, enhanced the binding of CAMSAP2 and 3 to microtubules and inhibited the rate of minus-end elongation compared to the effects of CAMSAP2 or CAMSAP3 alone **(Fig. 5H-N** and **Fig. S5D, E)**. No minus-end blocking was observed in these conditions. We conclude that WDR47 can be targeted to microtubules by all three CAMSAPs; it enhances the ability of CAMSAP1 to block minus-end outgrowth and the ability of CAMSAP2 and CAMSAP3 to accumulate on the microtubule lattice and slow down microtubule minus-end polymerization.

### WDR47 protects CAMSAP2/3-decorated microtubules against katanin-mediated severing

In the last set of experiments, we examined whether and how WDR47 can protect CAMSAP2/3-decorated microtubules from katanin-induced fragmentation. In the absence of ATP, 40 nM katanin p60/p80C strongly accumulated on the microtubule segments decorated by 40 nM CAMSAP3 without severing them **(Fig. 6A)**. The addition of 150 nM WDR47 significantly suppressed katanin binding **(Fig. 6A, B)**. The endogenous concentration of WDR47 was estimated to be 30-40 nM in HEK293 and Hela cells (Cho et al., 2022; Nagaraj et al., 2011), but could be higher in neurons, where this protein is most active. This indicates that WDR47 can compete with katanin for CAMSAP binding. In line with this idea, 150 nM WDR47 prevented amplification of microtubules decorated with 40 nM CAMSAP2 or CAMSAP3 by 40 nM p60/p80C and their destruction by 200 nM p60/p80C, allowing only limited microtubule fragmentation **(Fig. 6C-F, Fig. S6A** and **Video S5)**. WDR47 also prevented severing and subsequent amplification of CAMSAP3-decorated microtubules by 250 nM full-length katanin p60/p80 **(Fig. S6B** and **Video S5)**. Given that the binding sites of WDR47 and katanin on CAMSAP2 and CAMSAP3 are present next to each other (**Fig. S1A-C**), the ability of WDR47 to protect CAMSAP2/3-bound microtubules from katanin-mediated severing can be explained by the occlusion of the katanin-binding site on CAMSAP2 and CAMSAP3.

**Figure 6.**
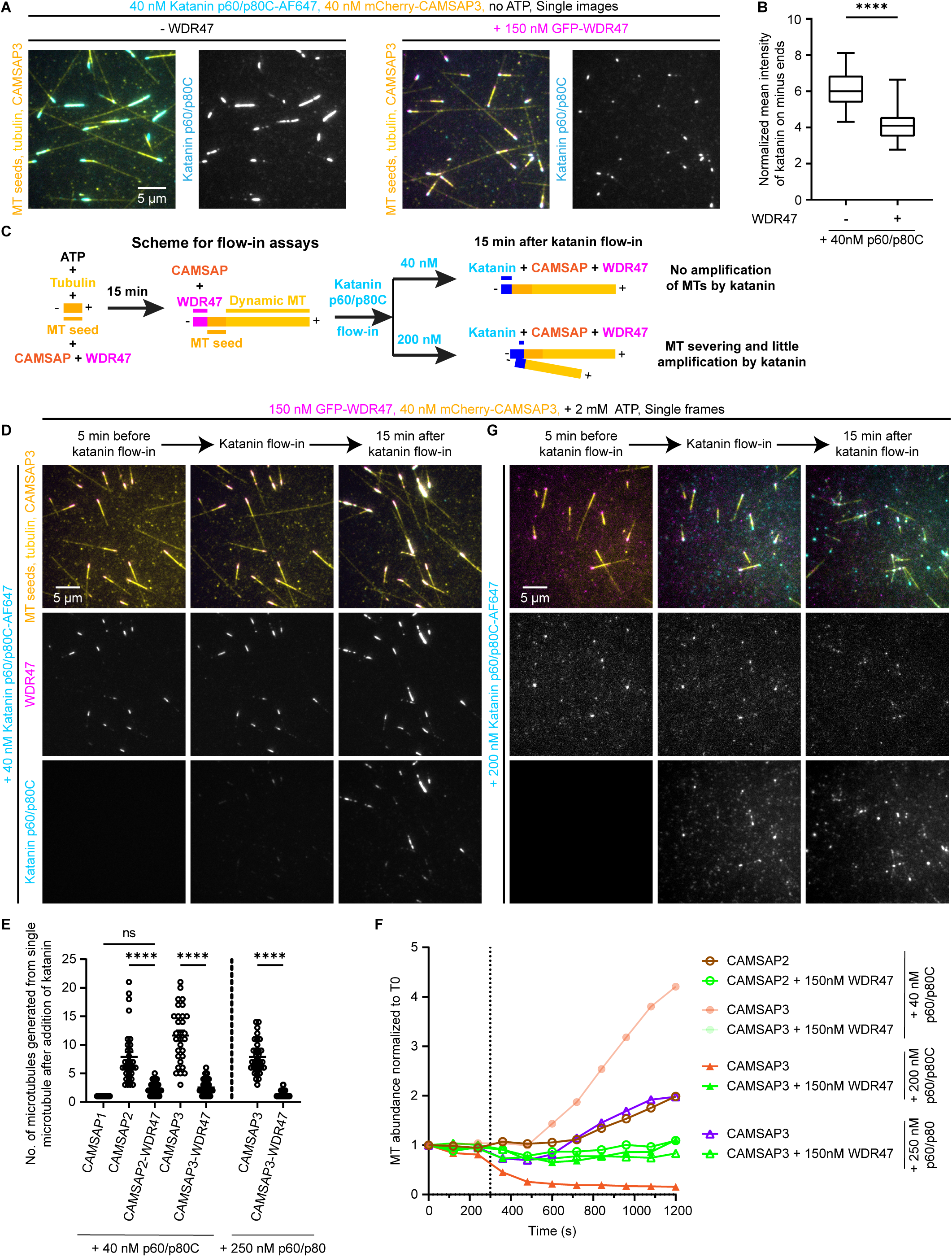
WDR47 protects CAMSAP-decorated microtubules against katanin-mediated severing. **(A)** Single images acquired from assays performed in two parallel chambers showing suppression of katanin (cyan) binding to CAMSAP3-decorated microtubule (yellow) minus-ends by WDR47 (magenta). These katanin assays were performed without addition of ATP to prevent severing activity of 40 nM katanin p60/p80C-SNAP-AF647 in the presence of 40 nM mCherry-CAMSAP3, 15 μM tubulin (left) or together with 150 nM GFP-WDR47 (right). **(B)** The box and whisker plot showing normalized mean intensity of katanin on CAMSAP3-decorated minus ends with or without WDR47, quantified (see method section for details) from the assays represented in panel **A**. n=31, N=3 for assays without GFP-WDR47; n=34, N=3 for assays with 150 nM GFP-WDR47, where n is the number of minus ends analyzed from N number of independent experiments. The box extends from the 25th to 75th percentiles, while the whiskers extend to the smallest and largest values. The line in the middle of the box is the median. Two-tailed unpaired t-test was used for significance test. ****p< 0.0001. **(C)** A scheme for in vitro reconstitution of microtubule severing by katanin and its regulation by CAMSAPs and WDR47. **(D, G)** Single frames (at indicated steps) from 20 min time-lapse movies showing WDR47 (magenta) binding to CAMSAP3-decorated microtubules (yellow) and a strong suppression in the microtubule severing and amplification by 40 nM katanin p60/p80C-SNAP-AF647 (cyan) **(D)** and in the microtubule destruction by 200 nM katanin p60/p80C-SNAP-AF647 (cyan) **(G)** in the presence of 150 nM GFP-WDR47, under indicated assay conditions, also depicted in panel **C**. See also (Fig. 2E, G). **(E)** Quantification of the number of microtubules generated from the minus-end of a CAMSAP-decorated microtubule as a result of katanin-mediated severing under the assay conditions represented in panels **(D)**, (Fig. 2E**), (Fig. S2B, D)**, (Fig. 3D) and **(Fig. S6A, B)**. n=49, N=3 for 10 nM CAMSAP1 and 40 nM p60/p80C; n=31, N=3 for 40 nM CAMSAP2 and 40 nM p60/p80C; n=52, N=3 for 40 nM CAMSAP2, 40 nM p60/p80C and 150 nM WDR47; n=30, N=3 for 40 nM CAMSAP3 and 40 nM p60/p80C; n=43, N=3 for 40 nM CAMSAP3, 40 nM p60/p80C and 150 nM WDR47; n=31, N=3 for 40 nM CAMSAP3 and 250 nM p60/p80; n=52, N=3 for 40 nM CAMSAP3, 250 nM p60/p80 and 150 nM WDR47; where n is the number of single microtubules analyzed from N number of independent assays. The plot presents mean ± s.e.m. and individual data points represent single microtubules analyzed. One-way ANOVA with Tukey’s multiple comparisons test was performed to compare the means with each other. ns= not significant, p= 0.0803; ****p< 0.0001. Data for 40 nM p60/p80C together with either CAMSAP1, or CAMSAP2, or CAMSAP3 are from Fig. 2F and data for 250 nM p60/p80 together with CAMSAP3 are from Fig. 3C, replotted here for direct comparison. **(F)** Quantification of microtubule abundance over time normalized to microtubule abundance at t=0 min (T0 being 5 min before flow-in) under the assay conditions indicated on plot, and represented in panels **(D, G)**, (Fig. 2E, G**), (Fig. S2B)** and **(Fig. S6A, B)**. n=3, N=3 for each condition, where n is the total number of single fields of view analyzed from N number of independent assays. Each individual curve represents the mean value of the three fields of view analyzed per condition. Data for 40 nM p60/p80C together with CAMSAP3 and 200 nM p60/p80C together with CAMSAP3 are from Fig. 2H; for 40 nM p60/p80C together with CAMSAP2 are from **(Fig. S2C)**; and for 250 nM p60/p80 together with CAMSAP3 are from Fig. 3E; replotted here for a direct comparison.

## Discussion

Microtubule organization and density are controlled by multiple factors with opposing activities, which can either stabilize microtubules or promote their disassembly. Here, we used in vitro reconstitution assays to dissect the regulatory mechanisms controlling the stability of free, non-centrosomal microtubule minus ends. We demonstrated that the microtubule severing enzyme katanin can counteract formation of long stretches of microtubule lattices stabilized by CAMSAP2 and CAMSAP3 in two ways: by inhibiting elongation of their minus ends and by cutting them into pieces. Interestingly, the combined activities of katanin and CAMSAPs at concentrations that are not far from the physiological can lead to strong microtubule amplification **(Fig. 7)**. Microtubule regulation by CAMSAP2 and 3 is conceptually simple – they decorate and stabilize microtubule lattices formed through minus-end polymerization (Jiang et al., 2014). The effects of microtubule severing enzymes, such as katanin, are more complex. These enzymes extract tubulin dimers from microtubule lattice causing their breakage and microtubule loss, but they can also amplify microtubules when the damaged microtubule lattices are repaired and serve as rescue sites (Vemu et al., 2018), or when the pieces of broken microtubules act as templates for microtubule re-growth (reviewed in (Kuo and Howard, 2021). In our in vitro assays, a combination of CAMSAP2/3 and katanin powerfully amplified microtubules, an effect that was stronger than previously observed with katanin alone or in combination with another microtubule minus-end-binding protein, ASPM (Jiang et al., 2017). This effect is likely caused by a combination of two factors. First, CAMSAP2 and CAMSAP3 can effectively stabilize freshly severed microtubules (Jiang et al., 2014). Second, katanin itself can bind and decorate the ends of microtubules it severs, because in addition to the ATPase domain, it also has a positively charged microtubule-binding domain which consists of the heterodimeric complex formed by α-helices in p60 N-terminus and p80 C-terminus and the linker region present in p60 (Jiang et al., 2017). Altogether, our in vitro assays suggest that in the absence of other cellular factors, a physiological concentration of CAMSAP2/3 and katanin would increase microtubule density **(Fig. 7)**.

**Figure 7.**
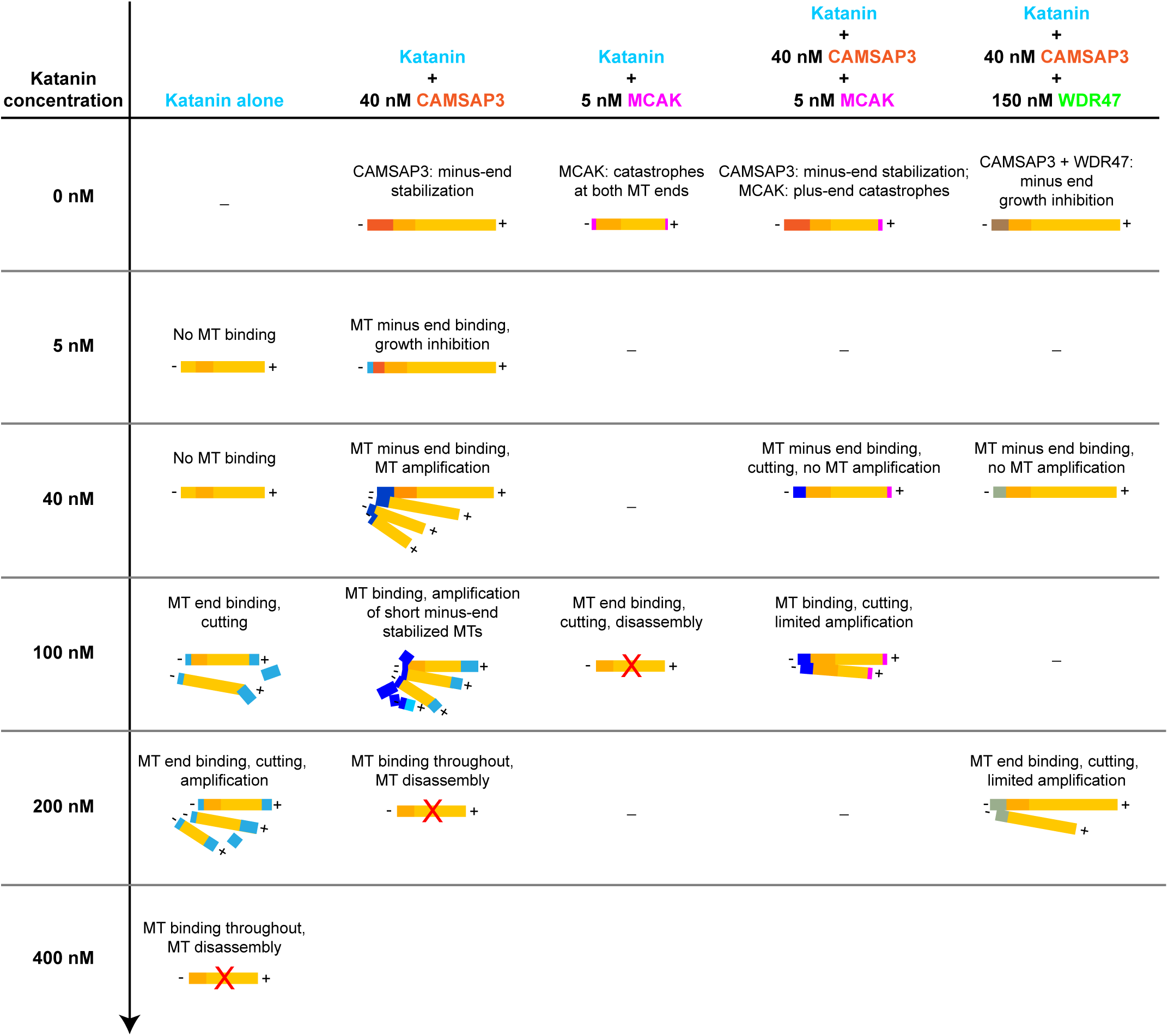
Katanin, CAMSAPs, kinesin-13 and WDR47 regulate microtubule abundance through their combined activities. Table summarizing the effects of different combinations of katanin, CAMSAPs, kinesin-13 and WDR47 on microtubule regulation.

This mechanism can be physiologically relevant: for example, the homologs of CAMSAPs and katanin cooperate to generate microtubules during neurite outgrowth in *C.elegans* (Shen et al., 2026).However, in other cellular settings, such as mammalian cancer cells and fibroblasts, there is no evidence of strong microtubule amplification caused by the combination of CAMSAPs and katanin (Jiang et al., 2018; Jiang et al., 2014). In fact, genetic screening showed that CAMSAPs and katanin have opposite effects on the formation of stabilized microtubules (Nieuwenhuis et al., 2017). Our in vitro experiments help to explain why this is the case. We found that kinesin-13, even at a concentration below physiological, very potently destroys the products of katanin-driven severing and strongly limits microtubule amplification by katanin even in the presence of a powerful microtubule stabilizer such as CAMSAP3 **(Fig. 7)**. Kinesin-13 family members do not walk on microtubules but rather peal off protofilaments at microtubule ends (Walczak et al., 2013). Since they act on individual protofilaments, they will efficiently disassemble not only new microtubule ends but also sites of microtubule lattice damage and repair, induced by a severing enzyme. Enhancement of spastin-induced microtubule severing by kinesin-13 KIF2A is fully in line with this idea (Henkin et al., 2023). Therefore, severing enzymes, even though they are not sufficiently abundant to disassemble microtubules and can by themselves even potentiate microtubule rescue and amplification (Vemu et al., 2018), ultimately often act as negative regulators of microtubule density, because they cooperate with microtubule depolymerases.

Our data show that the microtubule stability and numbers are very sensitive to the concentrations of katanin and kinesin-13. The destruction of CAMSAP-decorated microtubules by these factors is counterbalanced by WDR47, which seems to work by increasing CAMSAP accumulation on microtubules and at the same time preventing katanin binding to these microtubules **(Fig. 7)**. This is in agreement with studies in neurons, where the knockout or knockdown of WDR47 or increase in katanin activity led to a loss of CAMSAP2/3-decorated microtubules (Buijs et al., 2021; Chen et al., 2020). The finding that WDR47 helps to protect CAMSAP-decorated microtubule stretches is also in agreement with the fact that the loss of WDR47 can be partly compensated by overexpression of microtubule-binding fragments of CAMSAPs both in neurons and in multiciliated cells (Chen et al., 2020; Liu et al., 2021a). WDR47 and it cooperation with CAMSAP are conserved in worms, though in *C.elegans* neurons the homologue of WDR47 promotes rather than inhibits katanin-mediated CAMSAP-dependent microtubule severing (Shen et al., 2026).

Another interesting effect that has been uncovered by our in vitro reconstitutions is the cooperative inhibition of microtubule minus-end elongation by all studied factors. In vitro, both microtubule plus- and minus ends can grow, although plus-end polymerization occurs much faster. In cells, this difference is typically amplified further: microtubule plus-end growth is accelerated by specific factors such as XMAP215/chTOG (Brouhard et al., 2008; Howard and Hyman, 2007), whereas the minus ends are either capped by γ-TuRC, or their growth is inhibited by different factors. CAMSAP2 and particularly CAMSAP3 slow down tubulin addition to the minus-ends, possibly by affecting their structure (Atherton et al., 2017; Jiang et al., 2014). However, since growth of CAMSAP-decorated minus-ends is processive, one would still expect formation of long stretches of CAMSAP-bound microtubule lattices. One of the reasons why this does not happen might be further inhibition of minus-end growth by WDR47 and katanin. Restricting the length of the minus-end-grown and CAMSAP-decorated microtubule lattices by inhibiting minus-end polymerization would further reduce katanin-mediated severing and amplification because of fewer available cutting sites.

An interesting aspect of such regulation is the interplay between WDR47 and CAMSAP1. CAMSAP1 by itself does not decorate microtubules but rather tracks their minus ends. CAMSAP1 does not interact with katanin but it does strongly bind to WDR47, and together, they can efficiently block microtubule minus-end elongation. In cells, CAMSAP1 has microtubule-stabilizing effects in glia (Coquand et al., 2021), and its loss in neurons shows phenotypes similar to those of WDR47, such as formation of multiple axons and suppression of multipolar-bipolar transition during critical neuronal migration (Chen et al., 2020; Zhou et al., 2020). These data suggest that CAMSAP1 and WDR47 work together and that inhibition of microtubule minus-end dynamics is important for neuronal function.

To conclude, our multicomponent in vitro assays allowed us to dissect the interplay of opposing factors in controlling microtubule dynamics, stability and abundance. In future, it will be worthwhile to build their complexity even further, by including more proteins, for example microtubule stabilizers such as CLASPs, which seem to cooperate with CAMSAPs in minus-end stabilization (Wu et al., 2016), or by mimicking regulatory effects of protein modifications.

## Methods and Protocols

### DNA constructs

We used previously described katanin p60/p80C and katanin p60/p80 constructs (Jiang et al., 2017) and full-length MCAK construct (Atherton et al., 2017; Jiang et al., 2014). p80C-SNAP-Twin-Strep, katanin p80-SNAP-Twin-Strep and Twin-Strep-mCherry-MCAK constructs were made by cloning into a modified pTT5 expression vector (Addgene no. #44006) with a C-terminus SNAP-Twin-Strep tag or an N-terminus Twin-Strep-mCherry tag. GFP-WDR47 construct was described previously (Buijs et al., 2021). Twin-Strep-GFP-WDR47 was made by cloning the full-length construct in a modified C1 expression vector with an N-terminus Twin-Strep-GFP-tag. Twin-Strep-mCherry-CAMSAP1, Bio-Tev-mCherry-CAMSAP2, and Bio-Tev-mCherry-CAMSAP3 were made by cloning the previously described full-length constructs (Jiang et al., 2014) in modified C1 vectors with either a Twin-Strep-mCherry or Bio-Tev-mCherry tag at the N-terminus. Twin-Strep-tag has been abbreviated as SII-tag in the rest of the chapter.

### Cell lines and cell culture

HEK293T cells (from ATCC) were cultured in Dulbecco’s Modified Eagle’s Medium DMEM/Ham’s F10 media (1:1) supplemented with 10% fetal calf serum (FCS) and 1% antibiotics (penicillin and streptomycin) at 37 °C. This cell line was not found in the commonly misidentified cell lines database maintained by the Immunization Coalition of Los Angeles County. The cell lines were routinely checked for mycoplasma contamination using LT07-518 Mycoalert assay. Polyethylenimine (PEI, Polysciences) was used to transfect HEK293T cells with plasmids at 3:1 ratio of PEI:plasmid for StrepTactin- and Streptavidin-affinity based protein purification.

### Purification of recombinant proteins from HEK293T cells for in vitro reconstitution assays

Recombinant proteins used in the in vitro reconstitution assays (human GFP-WDR47, mCherry-CAMSAP1 and mCherry-MCAK, mouse p60/p80C-SNAP-AF647 and p60/p80-SNAP-AF647) used in the in vitro reconstitution assays were purified using Twin-Strep-tag and Strep-Tactin affinity purification method as described previously (Sharma et al., 2016). In brief, HEK293T cells were transfected with 50 μg of respective constructs per 15 cm dish and harvested 36 hours post-transfection from four 15 cm dishes each. For purification of katanin complex, 30 μg each of both the katanin subunits p60 and p80 constructs were transfected per 15 cm dish. Cells were resuspended and lysed in lysis buffer (50 mM HEPES, 300 mM NaCl, 0.5% Triton-X-100, 1 mM MgCl_2_ and 1 mM EGTA (also 1 mM ATP for katanin and MCAK), pH 7.4) supplemented with EDTA-free protease inhibitor cocktail (Roche). Cell lysates were clarified by centrifugation at 21,000xg for 20 min at 4°C. The supernatants obtained from the previous step were incubated with equilibrated Strep-Tactin Sepharose beads (28-9355-99, GE Healthcare) for 45 min at 4 °C. Following incubation, beads were washed three times with the wash buffer (50 mM HEPES, 300 mM NaCl and 0.1% Triton-X-100, 1 mM MgCl_2_, 1 mM EGTA and 1 mM DTT (also 1 mM ATP for katanin and MCAK), pH 7.4) and proteins were eluted for 15 min at 4 °C in elution buffer (50 mM HEPES, 150 mM NaCl (300 mM NaCl and also 1 mM ATP for katanin and MCAK), 0.05% Triton-X-100, 1 mM MgCl_2_, 1 mM EGTA, 1 mM DTT and 2.5 mM d-Desthiobiotin, pH 7.4).

For SNAP-tag labeling of katanin p80C or full-length p80 with Alexa Fluor 647 dye, washed beads were incubated with labeling mix (50 μM Alexa Fluor 647 dye in 50 mM HEPES, 150 mM NaCl and 0.1% Triton-X-100, 1 mM MgCl_2_, 1 mM EGTA, 1 mM ATP and 1 mM DTT, pH 7.4) for 1 hour. Following this incubation, beads were washed five times with wash buffer containing 300 mM NaCl to remove excess dye before elution. After elution, purified katanin was then subjected to buffer exchange using Vivaspin 500 centrifugal concentrator (10 kDa MWCO, Sartorius VS0102) for a final NaCl concentration of 150 mM in eluate.

For GFP-WDR47 and mCherry-MCAK purifications, following the incubation of the beads with the lysate, beads were additionally washed five times using high salt (1 M NaCl) containing wash buffer (50 mM HEPES, 0.1% Triton-X-100, 1 mM MgCl_2_ and 1 mM EGTA (also 1 mM ATP for katanin and MCAK), pH 7.4) before washing three times with 300 mM NaCl containing wash buffer.

For purification of human mCherry-CAMSAP2 and mouse mCherry-CAMSAP3, HEK293T cells were transfected with 25 μg of Bio-Tev-mCherry-CAMSAP2 and 25 μg of BirA per 15 cm dish. Cells were harvested 36 hours post-transfection from four 15 cm dishes and resuspended in lysis buffer (50 mM HEPES, 300 mM NaCl, 0.5% Triton-X-100, 1 mM MgCl_2_, 1 mM EGTA and 1 mM DTT, pH 7.4) supplemented with EDTA-free protease inhibitor cocktail (Roche). Cell lysates were incubated with Dynabeads M-280 streptavidin (Invitrogen-11206D) for 1 hour. Beads were washed thrice with lysis buffer without protease inhibitors and thrice with the TEV cleavage buffer (50 mM HEPES, 150 mM NaCl, 0.05% Triton-X-100, 1 mM MgCl_2_, 1 mM EGTA and 1 mM DTT). Proteins were eluted in 50 μl TEV cleavage buffer containing 0.5 μg of glutathione S-transferase-6x-histidine Tobacco etch virus protease site (Sigma-Aldrich) for 2 hr at 4 °C.

Purified proteins were immediately aliquoted, snap-frozen in liquid N2 and stored at -80 °C. The purity and composition of purified proteins were analyzed by Coomassie-staining of SDS-PAGE gels and mass spectrometry.

### Mass spectrometry

To assess the quality of purified recombinant proteins that were previously not characterized (GFP-WDR47 and katanin p60/p80-SNAP-AF647), samples were digested using S-TRAP micro filters (ProtiFi) according to the manufacturer’s protocol or were digested using the single-pot, solid-phase-enhance sample preparation method (SP3) (Hughes et al., 2019) protocol (SII-mCherry-MCAK). In brief, for the S-trap method, 4 μg of protein samples were denatured using 5% SDS buffer, then reduced and alkylated using DTT (20 mM, 10 min, 95 °C) and iodoacetamide (IAA, 40 mM, 30 min). Samples were subjected to acidification and precipitation using a methanol triethylammonium bicarbonate (TEAB) buffer before loading on a S-TRAP column. Trapped proteins were washed four times with methanol TEAB buffer for and then were digested overnight using Trypsin (1 μg, Promega) at 37 °C. For SP3 digestion of SII-mCherry MCAK, 2.2ug of protein was first reduced and alkylated by adding Tris(2-carboxyethyl)phosphine (TCEP) and chloroacetamide (CAA) to a final concentration of about 10 mM and 40 mM, respectively. Incubation first took place at 95 °C for 10 minutes, then in the dark for 20 minutes at room temperature (RT). Both hydrophilic and hydrophobic Cytiva Sera-Mag™ Carboxylate-Modified Magnetic Beads, beads were used in a 1:1 ratio, using 75ug beads in total. The beads were washed twice with 200 μl MQ water. 100% ethanol (EtOH) was added to the sample to a final concentration of 75% EtOH, and incubated in a Thermomixer at 1000 rpm for 20 minutes at RT. The tubes were placed in a magnetic rack and the supernatant was removed. The sample was washed two times with 80% EtOH, followed by a washing step with 100% acetonitrile (ACN). For digestion, the beads were resuspended in 200 μl 100 mM ammonium bicarbonate (AMBIC). Then, the samples were sonicated for two minutes in a water bath. The samples were digested using the proteases Trypsin (Promega) and Lys-C, in a 1:25 and 1:75 protease to protein ratio, respectively. Digestion was performed overnight in a Thermomixer at 1000 rpm, at 37 °C. The samples were spun down and acidified by adding 10% trifluoracetic acid (TFA) to a final concentration of 5% TFA. The beads were immobilized on the magnetic rack, and the acidified peptide solution was transferred to new Eppendorf tubes.

All digested peptide samples were eluted and dried in a vacuum centrifuge before LC-MS analysis. The samples were analysed using reversed phase nLC-MS/MS using an Ultimate 3000 UHPLC coupled to a Thermo Scientific^TM^ Orbitrap Exploris 480. For the GFP-WDR47 and katanin p60/p80-SNAP-AF647 samples, the digested peptides were separated over a 50 cm reversed phase column, packed in-house, (Agilent Poroshell EC-C18, 2.7 μm, 50 cm x 75 μm) using a linear gradient with buffer A (0.1% FA) and buffer B (80% acetonitrile, 0.1% FA) ranging from 13-44% B at a flow rate of 300 nL/min over 38 min, followed by a column wash and re-equilibration step, resulting in a total acquisition time of 55 min. For the SII-mCherry-MCAK sample, the digested peptides were dissolved in 2% formic acid (FA) ad separated on a 50-cm reversed-phase analytical column, with an integrated emitter, that has a 75 µm inner diameter and was packed in-house with ReproSilPur C18- AQ 1.9 µm resin (Dr. Maisch GmbH). Mobile Phase A and B were the same as mentioned above. The Acclaim Pepmap 100 C18 (5 mm × 0.3 mm, 5 μm, ThermoFisher Scientific) trap column was operated at a temperature of 32 °C, and the analytical column at a temperature of 50 °C. A flow rate of 0.3 μL/min was used. Starting at 4.0%, Buffer B was increased over a total gradient-time of 45 minutes using the following stepwise increases: 11% Buffer B (3-35 min), 30% Buffer B (35-40 min), and 44% Buffer B (40-44 min), and 55% Buffer B (44-45 min). The column was then washed out with 99% Buffer B for 5 minutes, followed by a 10-minute re-equilibration step with 4.0% Buffer B.

For all samples, the MS data was acquired using a DDA method with the set MS1 scan parameters in profile mode: 60,000 resolution, and a scan range of 375-1600 m/z. The automatic gain control (AGC) target was set equal to 3E6 and a maximum injection time of 20 ms for the two S-trap digested samples. The AGC and maximum injection time were set to ‘Auto’ and ‘Standard’, respectively, for the SII-mCherry-MCAK sample. The MS2 scan parameters were set at 15,000 resolution, with an AGC target set to standard, an automatic maximum injection time and an isolation window of 1.4 m/z. Scans were acquired first using a fixed mass of 120 m/z, a mass range of 200-2000 and a normalized collision energy (NCE) of 28. Precursor ions were selected for fragmentation using a 1 s scan cycle, a 10 s dynamic exclusion time and a precursor charge selection filter for ion possessing +2 to +6 charges.

The raw files for GFP-WDR47 and katanin p60/p80-SNAP-AF647 were processed using Proteome Discoverer (PD) (version 2.4, Thermo Scientific). A database search was performed using Sequest HT for MS/MS fragment spectra against a human database (UniProt, year 2020) that was modified to include protein sequences from our cloned constructs and a common contaminants database. A fragment mass tolerance of 0.06 Da and precursor mass tolerance of 20 ppm was set for the search parameters. Up to two missed cleavages were allowed by Trypsin digestion. Carbamidomethylation was set as fixed modification and methionine oxidation and protein N-term acetylation was set as variable modifications. Data filtering performed using percolator resulted in 1% FDR for peptide spectrum match (PSM) and a 1% FDR was applied to peptide and protein assemblies. An additional filter with a minimum Sequest score of 2.0 was set for PSM inclusion. MS1 based quantification was performed using the Pecursor Ion Quantifier node with default settings and precursor ion feature matching was enabled using the Feature Mapper node. Common protein contaminants from the results table were filtered out. The raw data of SII-mCherry-MCAK were analyzed using the MaxQuant-Andromeda softwar**e** (v.2.4.7.0) for protein identification, using the iBaq values for quantification. The MS/MS spectra were searched against the UniProtKB Human Proteome (organism_id: 9606, reviewed, canonical & isoform FASTA), and a separate FASTA file containing the recombinant protein. Default parameters were used for the precursor mass tolerance (20 p.p.m first search, 4.5 p.p.m. main search), and the False-Discovery Rate (FDR) was set to 1%. Up to three missed cleavages for both proteases were allowed. Carbamidomethylation of cysteine residues was set as a fixed modification, and oxidation of methionine and acetylation of the protein N-terminus were set as variable modifications, with a maximum of five modifications per peptide. The iBaq values from the MaxQuant search were imported into the Perseus software (v.2.0.11.0), where the protein groups were filtered out for ‘Reverse’ (false positives), ‘Contaminants’, and ‘Only identified by site’, as well as having only one peptide hit.

### In vitro reconstitution assays

#### Preparation of GMPCPP-stabilized microtubule seeds

Double-cycled GMPCPP-stabilized microtubules used for in vitro assays were prepared as described previously (Mohan et al., 2013). In brief, GMPCPP-stabilized microtubules seeds were prepared in the presence of GMPCPP (Guanylyl-(a,b)-methylene-diphosphonate (Jena Biosciences)) by two rounds of polymerization and a depolymerization cycle. First, a 20 μM porcine brain tubulin (Cytoskeleton) mix composed of 70% porcine unlabeled tubulin, 18% biotin tubulin and 12% labeled-tubulin (HiLyte488/rhodamine/HiLyte647-tubulin) was incubated with 1 mM GMPCPP in MRB80 buffer (pH 6.8, 80 mM K-PIPES, 1 mM EGTA and 4 mM MgCl_2_) for 30 minutes at 37 °C. Polymerization mix was then pelleted by centrifugation in an Airfuge at 119,000xg for 5 min followed by resuspension and depolymerization in MRB80 buffer for 20 min on ice and subsequent polymerization for 30 min in the presence of fresh 1 mM GMPCPP at 37 °C. GMPCPP-stabilized microtubule seeds were pelleted, resuspended in MRB80 buffer containing 10% glycerol, aliquoted, snap-frozen in liquid N2 and stored at -80 °C.

#### In vitro reconstitution of microtubule dynamics

In vitro reconstitution assays were performed as previously described (Mohan et al., 2013). Flow chamber was assembled using plasma-cleaned glass coverslip (18 x 18 mm coverslips) and microscopic slide attached together using double-sided tape. These chambers were then functionalized by 5 min incubation with 0.2 mg/ml PLL-PEG-biotin (Susos AG, Switzerland) followed by 5 min incubation with 1 mg/ml NeutrAvidin (Invitrogen) in MRB80 buffer. Next, biotinylated-GMPCPP-stabilized microtubule seeds were attached to the coverslip through biotin-NeutrAvidin links by incubating the chamber for 3 min. Non-immobilized seeds were washed away with MRB80 buffer and flow chambers were further incubated with 0.8 mg/ml k-casein to prevent non-specific protein binding. The reaction mix with or without proteins (MRB80 buffer supplemented with 15 μM porcine brain tubulin (14.5 μM unlabeled porcine tubulin and 0.5 μM HiLyte488/rhodamine/HiLyte647-tubulin), 80 mM KCl, 1 mM GTP, 0.5 mg/ml k-casein, 0.1% methylcellulose, and oxygen scavenger mix (50 mM glucose, 400 mg/ml glucose-oxidase, 200 mg/ml catalase, and 4 mM DTT)) were added to the flow chambers after centrifugation in an ultracentrifuge (Beckman Airfuge) for 5 minutes at 119,000xg. Concentrations of the proteins and composition of the reaction mix have been indicated in either figures or figure legends. The flow chambers were then sealed with high-vacuum silicone grease (Dow Corning), and three consecutive 10 min time-lapse movies of 3 different ROIs from the flow chambers were acquired after 2 min incubation (time, t=0) on total internal reflection fluorescence microscope stage at 30 °C. All tubulin products were from Cytoskeleton.

#### Flow-in assays for microtubule severing

For flow-in assays, special flow-in chambers were prepared using 24 x 32 mm coverslips attached orthogonally with the slides using double sided tape. Flow chamber was functionalized as mentioned above and GMPCPP-stabilized microtubule seeds were attached to the coverslip through biotin-neutravidin links. Flow chamber was then incubated with 0.8 mg/ml k-casein to prevent non-specific protein binding. Next, a microtubule polymerization mix (15 μM tubulin (14.5 μM unlabeled porcine tubulin and 0.5 μM HiLyte488/rhodamine/HiLyte647-tubulin), 80 mM KCl, 1 mM GTP, 2 mM ATP (in all the severing assays, unless specified otherwise **(Fig. 6A, B)**), 0.5 mg/ml k-casein, 0.1% methylcellulose, and oxygen scavenger mix (50 mM glucose, 400 mg/ml glucose-oxidase, 200 mg/ml catalase, and 4 mM DTT) in MRB80 buffer with or without proteins centrifuged in an Airfuge for 5 min at 119,000xg) was added to the chamber and a 10 min time-lapse movie was acquired after 2 min incubation (time, t=0) on TIRF microscope stage at 30 °C to monitor the polymerization of microtubule minus-ends. Immediately, another 20 min time-lapse movie was acquired during which the microtubule severing mix (polymerization mix containing katanin depending upon the condition) was flowed into the chamber at 5 min after start of this movie.

### Image acquisition, processing and data analysis

#### TIRF microscopy

In vitro assays were imaged on an iLas2 TIRF microscope setup as described previously (Sharma et al., 2016). Briefly, ILas2 system (Roper Scientific, Evry, France) is a dual laser illuminator for azimuthal spinning TIRF illumination. This system was installed on the Nikon Eclipse Ti-E inverted microscope with a perfect focus system. This microscope was equipped with Nikon Apo TIRF 100x 1.49 N.A. oil objective (Nikon), CCD camera CoolSNAP MYO M- USB-14-AC (Roper Scientific) and EMCCD Evolve mono FW DELTA 512x512 camera (Roper Scientific) with the intermediate lens 2.5X (Nikon C mount adaptor 2.5X), 150 mW 488 nm laser, 100 mW 561 nm laser and 49002 and 49008 Chroma filter sets and controlled with MetaMorph 7.10.2.240 software (Molecular Devices). The final magnification using Evolve EMCCD camera was 0.064 μm/pixel. Temperature was maintained at 30 °C to image the in vitro assays using a stage top incubator model INUBG2E-ZILCS (Tokai Hit). Time-lapse images were acquired on Photometrics Evolve 512 EMCCD camera (Roper Scientific) at 3 s time interval with 100 ms exposure time for 10 or 20 minutes.

#### Microtubule growth dynamics analysis

Images and movies were processed using Fiji (https://imagej.net/Fiji). Kymographs from the in vitro reconstitution assays were generated using the ImageJ plugin KymoResliceWide v.0.4 https://github.com/ekatrukha/KymoResliceWide. Microtubule dynamics parameters viz. plus-end growth rate and catastrophe frequency were determined from kymographs using an optimized version of the custom-made JAVA plugin for ImageJ as described previously (Gouveia et al., 2010; Mohan et al., 2013).

#### Quantification of the number of microtubules generated upon katanin-mediated severing

Microtubules that were generated after flow-in from katanin-mediated cutting/fragmentation of polymerized minus end lattice (decorated by CAMSAP2/3 or tracked by CAMSAP1) and then subsequent cutting of the minus ends of these newly generated microtubules were counted into the total number of microtubules generated from a parent single microtubule. Microtubules generated as a result of cutting at the polymerized plus end lattice or microtubule crossovers were not considered in this analysis.

#### Quantification of microtubule abundance in the flow-in assays

Microtubule abundance was quantified using the Fiji plugin FeatureJ: Hessian (https://imagescience.org/meijering/software/featurej/) with its source code available at https://github.com/imagescience/FeatureJ/. For quantifications, we used the images of tubulin channel from a 20 min time-lapse movie acquired at 3 s time interval. To enhance the filament line-like structures using FeatureJ in “Hessian” mode, we calculated images of the smallest eigenvalue of Hessian tensor using the smoothing scale of 2 pixels. Then we applied an intensity threshold (variable per image stack, according to the signal intensity) to this eigenvalue stack, separating the microtubule filaments signal from the background. The obtained binary stack was skeletonized using built-in Fiji function to find only the centerlines of the filaments and remove residual variability in curves thickness. To quantify cumulative filament length per frame, we used integrated density (integrated intensity of skeletonized image) values (IntD_t_) of a fixed-size square ROI (450x450 pixels, 28.8x28.8 μm^2^) drawn in the center of the field of view. IntD_t_ at frame *t* was normalized to integrated density value at the first frame (IntD_t0_) and plotted as I_t_ over time. Therefore, normalized cumulative filament length (microtubule abundance) at any time point *t* was equal to I_t_ = IntD_t_ / IntD_t0_.

#### Analysis of minus-end growth behaviors

All the microtubule minus-end growth events were categorized into 3 categories: no blocking-when the minus ends started growing from the start of the movie; temporarily blocked- when minus were blocked for a short time from the beginning of the movie and then transitioned to a growing phase; completely blocked- when minus ends were not growing at all and remained totally blocked for the entire 10 min duration of the movie. If the minus end did not grow for 20 frames during a 10 min duration of the movie acquired at 3 s time interval, it was considered a block in the growth. The slowest minus-end growth rate observed across all the growth events analyzed was 0.07 μm/min.

#### Quantification of normalized mean intensity of CAMSAPs and katanin on minus ends

Two parallel flow chambers were assembled on the same plasma-cleaned glass coverslip for one experiment and microtubule dynamics assays were reconstituted as described above. CAMSAP2 **(Fig. 5L)** or CAMAP3 **(Fig. 5M)** in vitro reconstitutions with or without 150 nM GFP-WDR47 were performed in the parallel chambers. Samples were focused first on one area and still images of unexposed coverslip regions (ROIs) were acquired for each chamber after 10 min incubation time using the same acquisition settings and EMCCD camera on the TIRF microscope as described above. Microtubule minus ends were traced using a 10-pixels wide straight line, and the length and mean intensities of CAMSAP2/3 stretches were calculated. Similarly, mean intensities of CAMSAP2/3 on GMPCPP seeds were also calculated from the same image and its average was calculated from three GMPCPP seeds. Then, mean intensity of CAMSAP2/3 on each minus end was normalized to the average of that on GMPCPP seed from that assay. These normalized intensities for CAMSAP2/3 with or without WDR47 were finally plotted from three such independent experiments.

Normalized mean intensity of katanin on minus ends with or without WDR47 **(Fig. 6B)** was also quantified in the similar manner as described for CAMSAPs.

### Statistical analysis

All statistical details of experiments including the definitions, exact values of number of measurements, number of replicates, precision measures and statistical tests performed are mentioned in the figure legends. Data processing and statistical analysis were done in Excel and GraphPad Prism 10.4.2. Significance was defined as: ns = not significant for p > 0.05, *p < 0.05, **p < 0.01, ***p < 0.001 and ****p < 0.0001.

## Data availability

All data that support the conclusions are either available in the manuscript itself or available from the authors on request.

## Code availability

ImageJ macros used in this study are either available online at https://github.com/ekatrukha/KymoResliceWide, https://github.com/imagescience/FeatureJ, or from the corresponding authors on request.

## Acknowledgements

This work was supported by the European Research Council Synergy grant 609822 and a grant from the Swiss National Science Foundation 310030M_215014 to A.A. The authors acknowledge support received from the Dutch Research Council-funded Netherlands Proteomics Centre through the National Roadmap for Large-scale Research Infrastructures programme X-Omics (project 184.034.019) to K.S. for the mass spectrometry analysis.

## Supplemental figure legends

**Figure S1, related to Figure 1.**
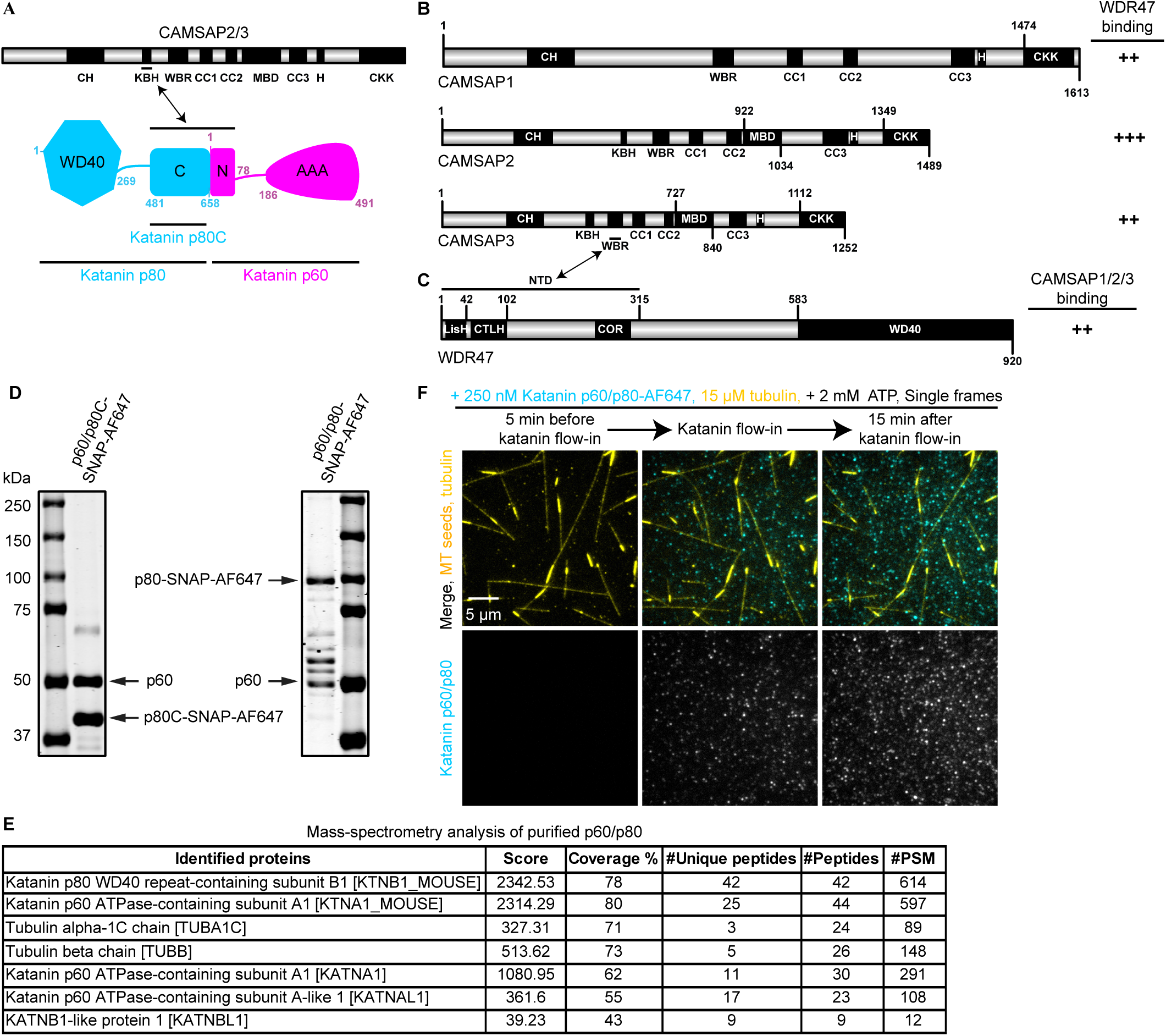
Schematic representation of the domain structure and interactions of katanin, CAMSAPs and WDR47, and characterization of purified full-length katanin. **(A-C)** Schematic representation of the domain structure of katanin **(A)**, CAMSAP proteins **(B)**, WDR47 **(C)**, and a summary of their known interactions. Indicated domains of katanin: AAA, ATPases associated with diverse cellular activities domain of p60; C, conserved C-terminal domain of p80; N, N-terminal microtubule-interacting and -trafficking domain of p60; WD40, WD40 repeat domain. Indicated domains of CAMSAPs: CC, coiled coil; CH, calponin-homology domain; CKK, CKK domain; H, alpha-helix; KBH, katanin-binding helix; MBD, microtubule-binding domain; WBR, WDR47-binding region. Indicated domains of WDR47: COR, cross-over region; CTLH, C-terminal to LisH domain; LisH, lissencephaly-1 homology domain; WD40, WD40 repeat domain. Protein schemes were generated using illustrator of biological sequences (IBS) software (Liu et al., 2015). **(D)** Coomassie-stained SDS-PAGE gels loaded with purified katanin p60/p80C-SNAP-AF647-SII and full-length katanin p60/p80-SNAP-AF647-SII. **(E)** List of proteins identified by mass spectrometry that were co-purified with full-length katanin p60/p80-SNAP-AF647-SII showing absence of any microtubule associated proteins in top hits. **(F)** Single frames (at indicated steps) from a 20 min time-lapse movie showing lack of katanin (cyan) localization to microtubules (yellow) and, thereby, lack of severing by 250 nM full-length katanin p60/p80-SNAP-AF647 alone. Rest of the experimental conditions are the same as shown for katanin p60/p80C-SNAP-AF647 in **Fig. 1A**. Assays were independently replicated three times.

**Figure S2, related to Figure 2.**
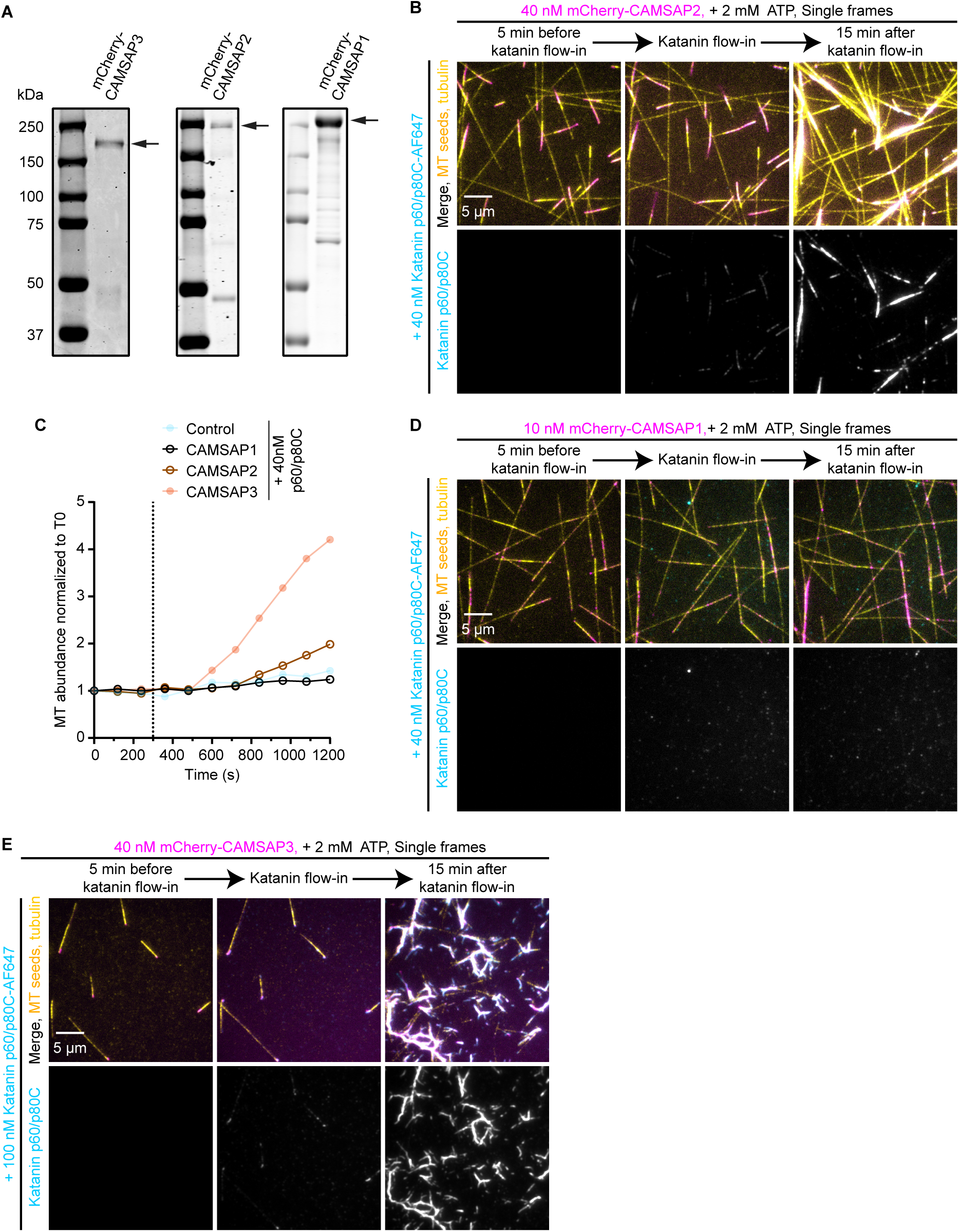
Characterization of the effects of katanin-CAMSAP combinations on microtubule abundance. **(A)** Coomassie-stained SDS-PAGE gels loaded with purified mCherry-CAMSAP3, mCherry-CAMSAP2 or mCherry-CAMSAP1. **(B)** Single frames (at indicated steps) from a 20 min time-lapse movie showing katanin p60/p80C-SNAP-AF647 (cyan) mediated severing and amplification of microtubules (yellow) decorated with mCherry-CAMSAP2 (magenta). **(C)** Quantification of microtubule abundance over time normalized to microtubule abundance at t=0 min (T0 being 5 min before katanin flow-in) in the presence of 40 nM katanin p60/p80C-SNAP-AF647 either alone or together with mCherry-CAMSAP1, mCherry-CAMSAP2, or mCherry-CAMSAP3 under the experimental conditions represented in panels **(B, D)**, **Fig. 1B** and **Fig. 2E**. n=3, N=3 for each condition, where n is the total number of single fields of view analyzed from N number of independent assays. Each individual curve represents the mean value of the three fields of view analyzed per condition. Data for 40 nM katanin p60/p80C alone and together with 40 nM CAMSAP3 are from **Fig. 1F** and **Fig. 2H** respectively, replotted here for comparison. **(D)** Single frames (at indicated steps) from a 20 min time-lapse movie showing lack of katanin (cyan) localization to mCherry-CAMSAP1-bound (magenta) microtubules (yellow) and, thereby, lack of severing by 40 nM katanin p60/p80C-SNAP-AF647. Experimental conditions in **(B, D)** are the same as shown for mCherry-CAMSAP3 in **Fig. 2E**. **(E)** Single frames (at indicated steps) from a 20 min time-lapse movie showing severing and amplification of mCherry-CAMSAP3-decorated (magenta) microtubules (yellow) by 100 nM katanin p60/p80C-SNAP-AF647 (cyan). Rest of the assay conditions are the same as shown for different katanin concentrations in **Fig. 2B, E, G**.

**Figure S3, related to Figure 3.**
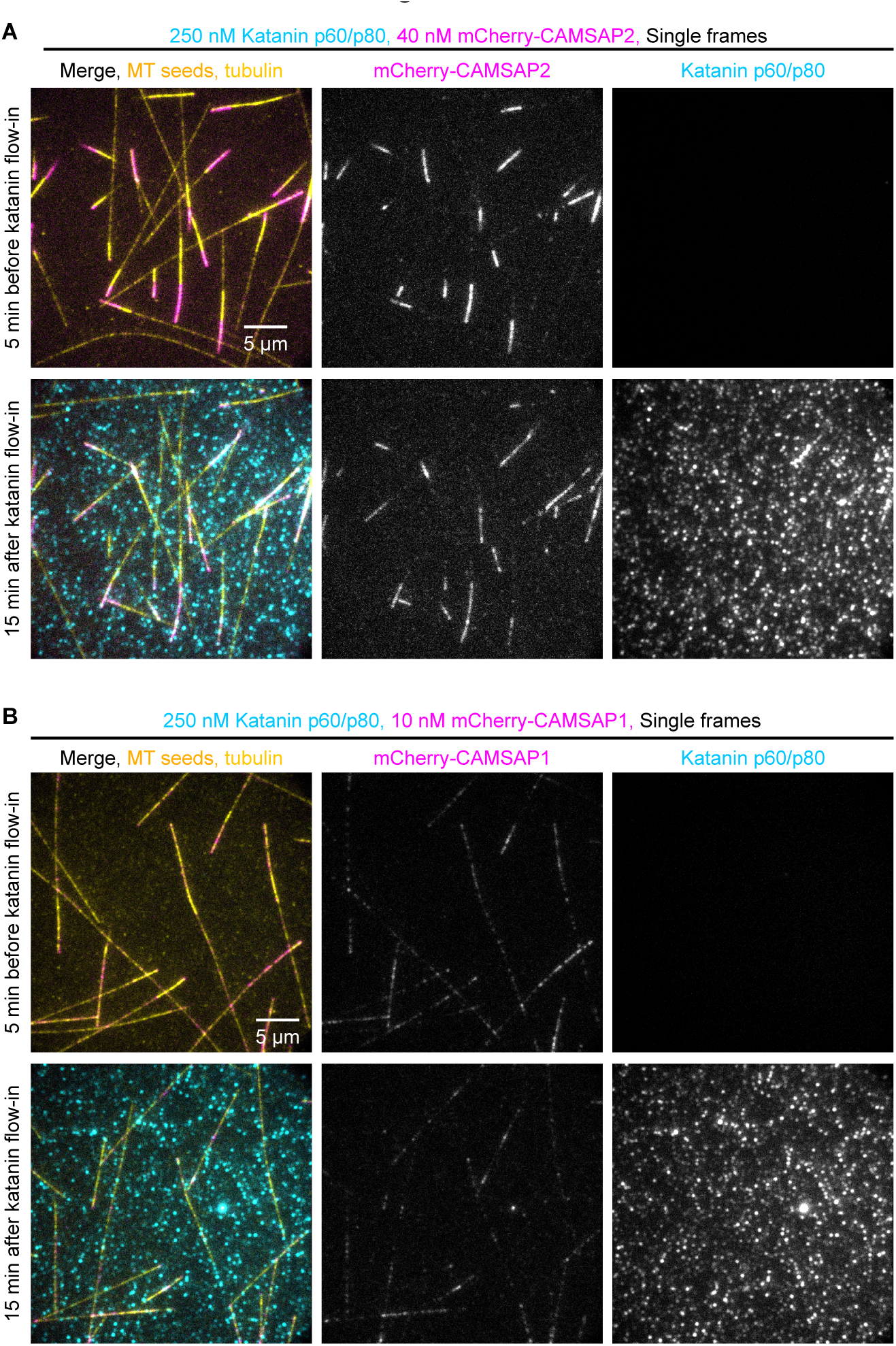
Characterization of the effects of full-length katanin and CAMSAPs combinations for microtubule severing experiments. **(A, B)** Single frames (at indicated steps) from 20 min time-lapse movies showing lack of any significant katanin (cyan) localization to microtubules (yellow) decorated with either 40 nM mCherry-CAMSAP2 (magenta) in **(A)**, or 10 nM mCherry-CAMSAP1 (magenta) in **(B)**, thereby, lack of severing by 250 nM full-length katanin p60/p80-SNAP-AF647 (cyan). Rest of the assay conditions are the same as shown for 40 nM mCherry-CAMSAP3 in **Fig. 3D**.

**Figure S4, related to Figure 4.**
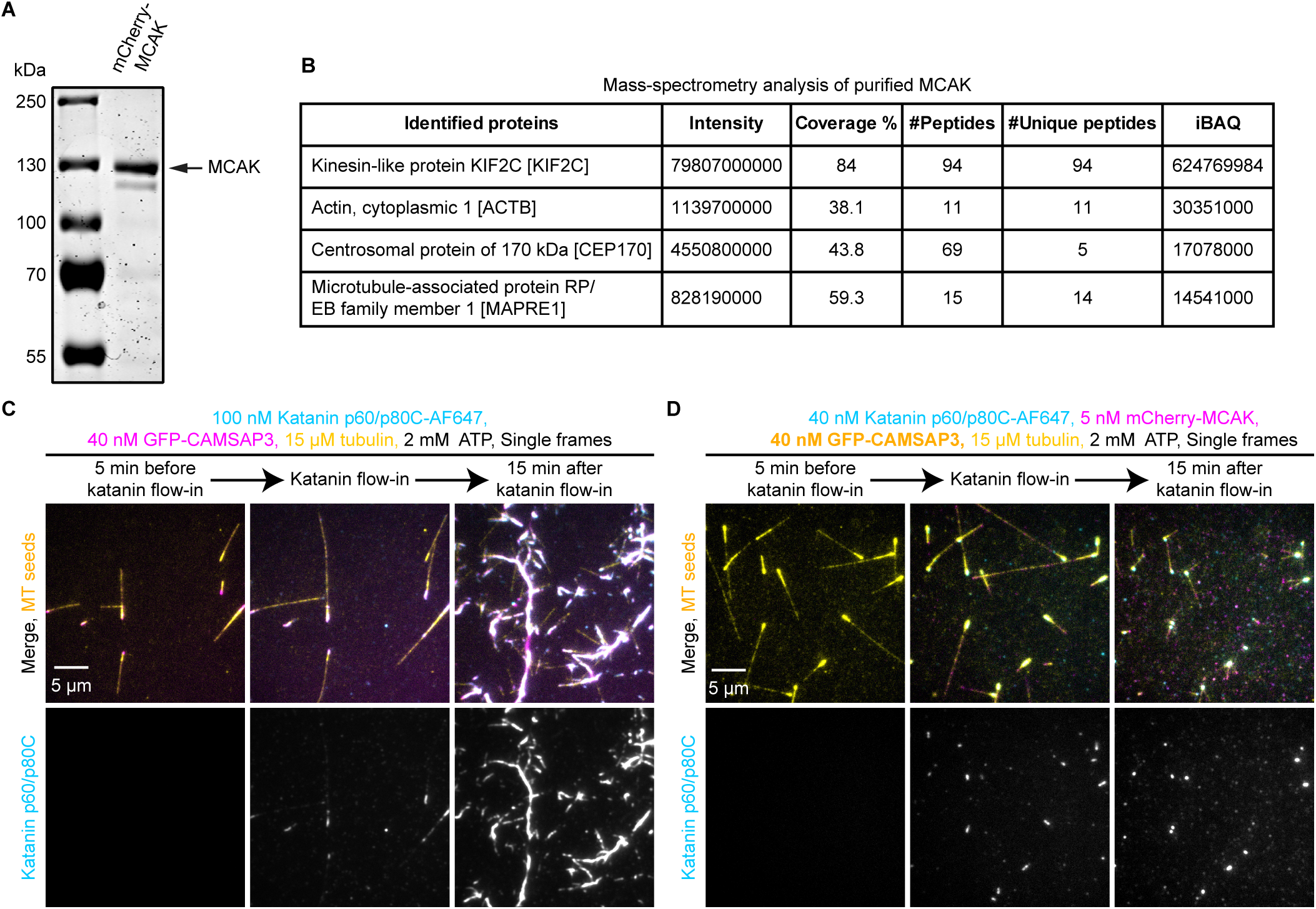
Characterization of purified MCAK and effective concentrations of katanin, CAMSAP3 and MCAK for microtubule severing experiments. **(A)** Coomassie-stained SDS-PAGE gels loaded with purified mCherry-MCAK. **(B)** List of proteins identified by mass spectrometry that were co-purified with SII-mCherry-MCAK. Note the mild contamination by CEP170 (2.7%) MAPRE1 (2.3%), calculated by taking their respective iBAQ ratios, in top hits. **(C)** Single frames (at indicated steps) from a 20 min time-lapse movie showing severing mediated amplification of microtubules (yellow) decorated with 40 nM GFP-CAMSAP3 (magenta) by 100 nM katanin p60/p80C-SNAP-AF647 (cyan), under indicated assay conditions, also depicted in **Fig. 4F**. **(D)** Single frames (at indicated steps) from a 20 min time-lapse movie showing steady-state maintenance of microtubule (yellow) abundance by the antagonizing activity of 5 nM mCherry-MCAK (magenta) against microtubule amplification by the combined activities of 40 nM GFP-CAMSAP3 (yellow) and 40 nM katanin p60/p80C-SNAP-AF647 (cyan) under indicated assay conditions, also depicted in **Fig. 4F**.

**Figure S5, related to Figure 5.**
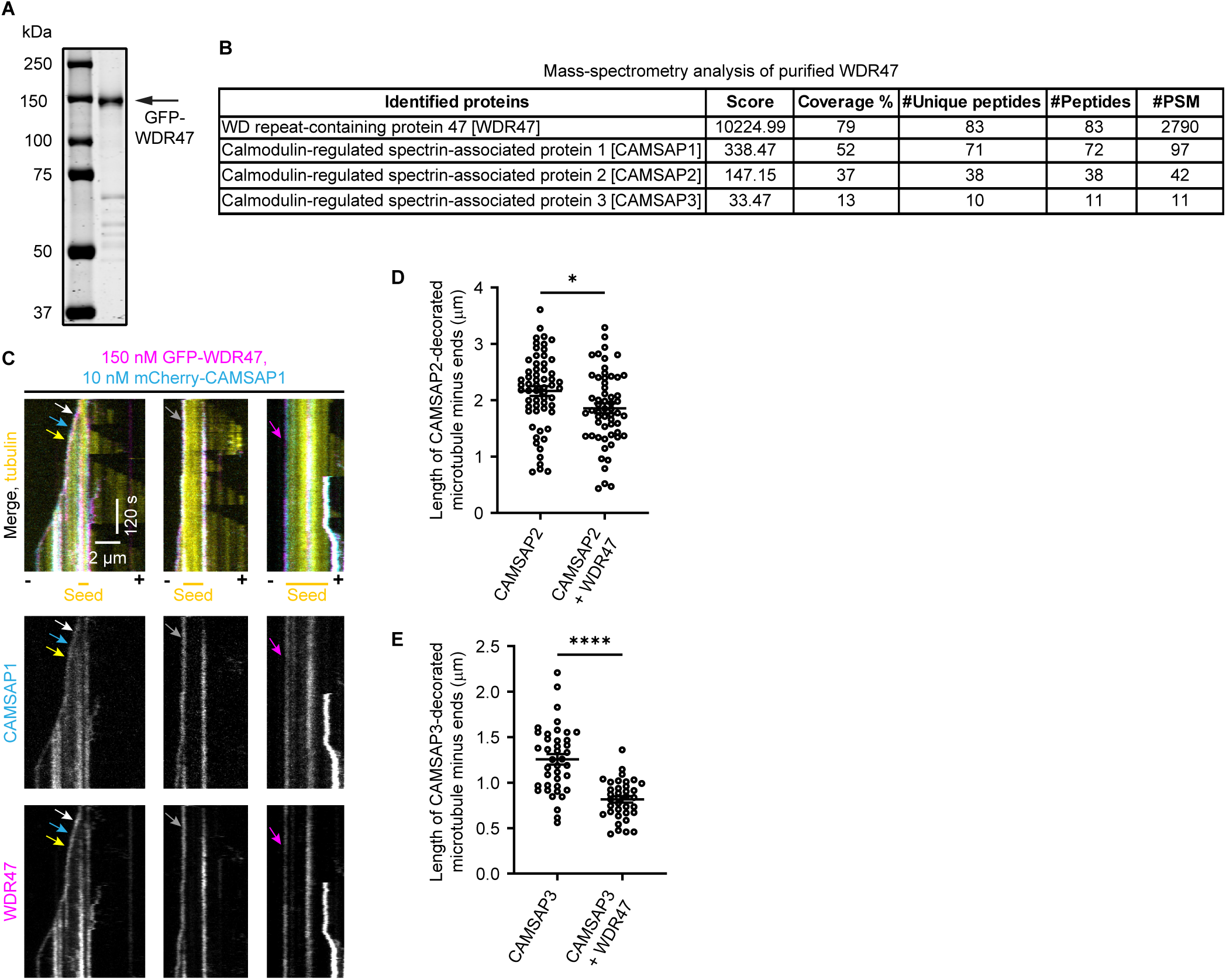
Characterization of purified WDR47 and effects of its binding interaction with CAMSAPs on microtubule mins-end dynamics. **(A)** Coomassie-stained SDS-PAGE gel loaded with purified GFP-WDR47. **(B)** List of proteins identified by mass spectrometry shows presence of all the three CAMSAPs that were co-purified with GFP-WDR47. **(C)** Representative kymographs from in vitro reconstitutions of microtubule (yellow) dynamics in the presence of 15 μM tubulin (yellow), 10 nM mCherry-CAMSAP1 (cyan) and 150 nM GFP-WDR47 (magenta). Three different types of minus-end growth behaviors are illustrated (also depicted in the key for figure panel **(Fig. 5D**): minus-end growth (left and yellow arrows); transition from temporarily blocked minus-end outgrowth to growing minus-end (middle and gray arrows) and completely blocked minus-end of a GMPCPP-stabilized microtubule seed (right and magenta arrows). White and cyan arrows denote tracking of minus ends by WDR47 and CAMSAP1, respectively. **(D, E)** Plots showing length of CAMSAP2 **(D)** and CAMSAP3 **(E)** stretches quantified from the assay conditions indicated on the plots and represented in **(Fig. 5H-K)**. In panel **D**:15 μM tubulin together with either 40 nM CAMSAP2 (n=60, N=3) or 40 nM CAMSAP2 and 150 nM GFP-WDR47 (n=57, N=3). In panel **E**:15 μM tubulin together with either 40 nM CAMSAP3 (n=40, N=3) or 40 nM CAMSAP3 and 150 nM GFP-WDR47 (n=36, N=3). n is the number of minus ends analyzed, and N is the number of independent experiments analyzed for each condition. The plot presents mean ± s.e.m. and individual data points represent each minus end analyzed. Two-tailed unpaired t-test was used to test for significance. *p= 0.0128; ****p< 0.0001.

**Figure S6, related to Figure 6.**
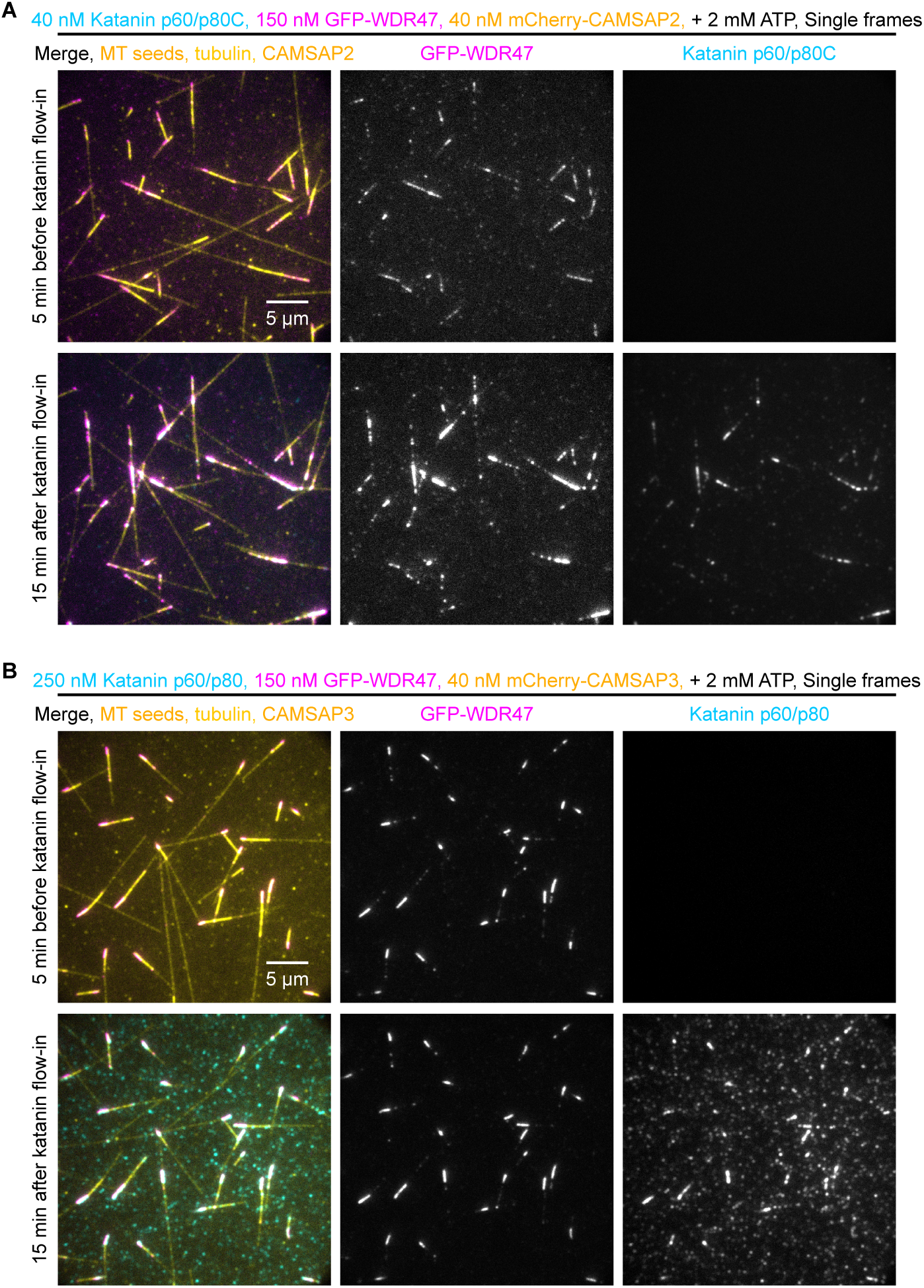
WDR47 prevents amplification of CAMSAP2/3-decorated microtubules against katanin-mediated severing. **(A, B)** Single frames (at indicated steps) from 20 min time-lapse movies showing WDR47 (magenta) binding to CAMSAP-decorated microtubules (yellow) and a strong suppression in microtubule severing and amplification by katanin (cyan) under following assay conditions: 40 nM mCherry-CAMSAP2, 40 nM katanin p60/p80C-SNAP-AF647 and 150 nM GFP-WDR47 **(A)** and 40 nM mCherry-CAMSAP3, 250 nM katanin p60/p80-SNAP-AF647 and 150 nM GFP-WDR47 **(B)**, under indicated assay conditions, also depicted in panel **6C**. See also **(Fig. S2B** and **Fig. 3D)**.

## Supplemental table legends

**Table S1, related to Figure S1 - Mass-spectrometry analysis of purified full-length katanin p60/p80**. List of proteins that were co-purified with full-length katanin p60/p80-SNAP-AF647-SII.

**Table S2, related to Figure S4 - Mass-spectrometry analysis of purified MCAK**. List of proteins that were co-purified with SII-mCherry-MCAK.

**Table S3, related to Figure S5 - Mass-spectrometry analysis of purified WDR47**. List of proteins that were co-purified with SII-GFP-WDR47.

## Supplemental video legends

**Video S1, related to Figure 1 -** Videos of 20 min time-lapse movies showing the absence or presence of microtubule severing, amplification and destruction by katanin p60/p80C-SNAP-AF647 at indicated concentrations in the video, ranging from low to very high: 40 nM, 100 nM, 200 nM, and 400 nM. Time-lapse images were acquired using TIRF microscope at 3 s time interval, constituting 401 frames and displayed at 50 fps.

**Video S2, related to Figure 2 -** Videos of 20 min time-lapse movies showing regulation of minus end dynamics, severing mediated amplification or complete disassembly of mCherry-CAMSAP3-decorated microtubules by 5 nM, 40 nM, 100 nM, and 200 nM katanin p60/p80C-SNAP-AF647, as indicated in the video. Time-lapse images were acquired using TIRF microscope at 3 s time interval, constituting 401 frames and displayed at 50 fps.

**Video S3, related to Figure 3 -** Video of a 20 min time-lapse movie showing severing mediated amplification of mCherry-CAMSAP3-decorated microtubules by 250 nM full-length katanin p60/p80-SNAP-AF647. Time-lapse images were acquired using TIRF microscope at 3 s time interval, constituting 401 frames and displayed at 50 fps.

**Video S4, related to Figure 4 -** Videos of 20 min time-lapse movies showing regulation of microtubule abundance by MCAK, katanin and CAMSAP3 under different conditions indicated in the video: depolymerization of microtubule ends by 5 nM mCherry-MCAK alone; microtubule destruction by 5 nM mCherry-MCAK and 100 nM katanin p60/p80C-SNAP-AF647; microtubule minus-end stabilization by 40 nM GFP-CAMSAP3 against depolymerization by 5 nM mCherry-MCAK; protection by 40 nM GFP-CAMSAP3 against microtubule destruction by 5 nM mCherry-MCAK and 100 nM katanin p60/p80C-SNAP-AF647. Time-lapse images were acquired using TIRF microscope at 3 s time interval, constituting 401 frames and displayed at 50 fps.

**Video S5, related to Figure 6 and Figure S6 -** Videos of 20 min time-lapse movies showing a strong suppression in the microtubule severing and amplification by 40 nM katanin p60/p80C-SNAP-AF647 or 250 nM full-length katanin p60/p80-SNAP-AF647, as well as in the microtubule destruction by 200 nM katanin p60/p80C-SNAP-AF647 in the presence of 150 nM GFP-WDR47. Time-lapse images were acquired using TIRF microscope at 3 s time interval, constituting 401 frames and displayed at 50 fps.

